# Active and total rhizosphere microbiomes of plant mixtures diverge from additive expectations, depending on nitrogen and plant composition

**DOI:** 10.64898/2026.09.04.749479

**Authors:** Jennifer E. Harris, Emma Rice, Carolyn Lowry, Estelle Couradeau, Liana T. Burghardt

## Abstract

Mixing plant functional groups can boost plant productivity, but the underlying rhizosphere microbial dynamics remain unclear. We tested when to expect divergence from additive expectations by measuring plant traits and active and total rhizosphere microbiomes (via BONCAT-FACS and 16S rRNA gene amplicon sequencing) in monocultures and mixtures of legumes, brassicas, and grasses under varying nitrogen. While shoot biomass was predictable from monocultures, legume-containing mixtures yielded greater-than-expected root biomass and N-fixation. Each plant species supported distinct active and total microbiomes, and nitrogen addition increased microbiome divergence among plant species and their mixtures. In most plant mixtures, microbiomes were additive combinations of those associated with the individual species; legume–grass combinations were a notable exception, showing non-additive shifts in microbial composition, microbial activity, and the abundance of Rhizobium sp. This divergence from additive expectations coincided with non-additive biomass gains in legume-grass mixtures, suggesting that emergent plant-microbe interactions may underlie synergistic belowground responses.

## Introduction

Increasing plant functional diversity can increase plant productivity (Hector *et al*. 1999; Hooper *et al*. 2005; Hooper & Dukes 2004; Mueller *et al*. 2013; Tilman *et al*. 1997). Plant mixtures can exhibit non-additive effects, such as overyielding, in which productivity exceeds that expected from monocultures (De Witt 1960). However, plant mixtures do not always produce non-additive effects (Dee *et al*. 2023; Urgoiti *et al*. 2022), so it is important to understand the mechanisms underlying them. Several mechanisms may contribute to non-additive effects in mixtures, such as competition, complementary, or facilitation. In niche complementarity, different plant species partition space and resources, thereby increasing productivity in a given area (Hooper *et al*. 2005). Under facilitation, one species may ameliorate harsh conditions, allowing other species to thrive (Bertness & Callaway 1994). Alternatively, increased plant productivity may depend on the species present rather than on diversity (Huston 1997). Soil microorganisms can be important mediators of these mechanisms, thereby influencing the benefits derived from plant functional diversity. A current gap in the implementation of plant mixtures is the lack of understanding of microbial dynamics when plant functional groups are combined. Here, we address this gap by measuring shifts in the total and active rhizosphere microbiomes and in resulting plant traits when plants grow alone or in combination.

The soil microbiome can increase plant productivity in plant mixtures by reducing pathogenic microbes or by increasing beneficial microbes. Modeling demonstrates that soil pathogens can have density-dependent effects, in which disease prevalence increases when plants of the same species are planted together (Bever *et al*. 1997). Plant mixtures can ameliorate these effects by reducing contact among same-species plants (Schnitzer *et al*. 2011). Additionally, soil microbes may increase nutrient availability, thereby reducing competition in mixtures (Rodríguez *et al*. 2006; Schimel & Bennett 2004; Xie *et al*. 2022). For instance, the positive effects of plant diversity are limited in the absence of soil microbiota (Schnitzer *et al*. 2011). Furthermore, plants were more productive when grown in soil preconditioned by other plant species rather than by conspecifics (Hendriks *et al*. 2013). In corn, intercropping with other crops increased microbial and metabolite diversity, potentially facilitating beneficial interactions (Jiang *et al*. 2024). However, there could be trade-offs in the soil microbiota promoted by plants in mixtures, where an organism beneficial to one plant is suppressed by another. For example, brassica species produce secondary metabolites called glucosinolates, which exhibit antimicrobial activity in vitro (Aires *et al*. 2009) and may thus suppress both beneficial bacteria and pathogens. A reduction in beneficial rhizobia, which form associations with legumes and convert atmospheric nitrogen into a plant-accessible form, can reduce plant biomass (Batstone *et al*. 2023). In sum, there are many potential pathways by which the soil microbiome can influence outcomes in plant mixtures.

New methods enable evaluation not only of the total soil microbiome but also its active subset. This is a key advance, as more than 90% of bulk soil microbes are dormant (Blagodatskaya & Kuzyakov 2013). Dormant microbes may obscure patterns in the active microbial community, as even when plants are present, activity in the bulk soil can be as low as 1% (Harris *et al*. 2025). Furthermore, extracellular DNA from dead microorganisms, also called relic DNA, can obscure shifts in the microbial community (Carini *et al*. 2016). Bioorthogonal non-canonical amino acid tagging - fluorescence-activated cell sorting (BONCAT-FACS) has become an effective tool for probing the active microbial community in soils (Couradeau *et al*. 2019). Here, we measure the effects of plant mixtures on both total and active rhizosphere microbial communities and evaluate when and if they differ.

Cover crops are an ideal system for exploring microbial communities in plant mixtures. The use of cover crops, or plants grown primarily for their environmental benefits rather than for market sale, has increased in recent years to address challenges such as soil and nutrient loss in agroecosystems (Bowman *et al*. 2025; Groff 2015). Different cover crop species provide different benefits. For example, legumes increase plant-available nitrogen through their mutualism with nitrogen-fixing rhizobia, grasses promote soil carbon storage due to their high C: N ratio, and brassicas suppress pests through weed competition and allelopathy (Haramoto & Gallandt 2004; Kou *et al*. 2012; Singh *et al*. 2024; Snapp *et al*. 2005). Many of these benefits are mediated at least in part by microbial communities (Nair & Ngouajio 2012). Farmers grow mixtures of cover crops, hoping to combine the benefits of each functional group, and indeed, some studies find greater benefits in diverse cover crop mixtures, including increased weed suppression, nitrogen retention, and aboveground biomass (Blesh 2018; Finney & Kaye 2017). However, cover crop benefits can be context-dependent (Blanco-Canqui 2024; Florence & McGuire 2020). A meta-analysis of cover crop mixtures with at least three species found that such mixtures typically perform similarly to monocultures in biomass, weed suppression, and other services (Florence & McGuire 2020). Perhaps cover crop mixtures could be tailored to enhance beneficial microbes and improve performance relative to monocultures.

While it is well established that individual plant identity shapes soil microbiome composition (Cloutier *et al*. 2020; Finney *et al*. 2017; Richards *et al*. 2026), how multi-species combinations influence these communities—and deviate from monocultures—remains poorly understood. To investigate this, we conducted a greenhouse experiment in field soil of plant monocultures, two-way mixtures, and three-way mixtures of three plant functional groups: a legume (crimson clover, *Trifolium incarnatum*), a brassica (canola, *Brassica napus*), and a grass (triticale, *× Triticosecale*), under both ambient and nitrogen-amended conditions (Figure 1). We quantified plant outcomes alongside microbial activity level and the community structure of total, active, and inactive rhizosphere microbial communities. Non-additive effects were assessed by comparing observed mixture dynamics with additive expectations calculated as a weighted mean of sowing proportions or of observed relative biomass (De Witt 1960; Fowler 1982). Ultimately, we tested whether plant mixtures function as additive intermediates of their corresponding monocultures across plant traits, total microbiomes, and active microbiomes by addressing four central questions: 1) How do plant traits shift between monocultures and mixtures? 2) Do total microbial composition and diversity differ between these treatments, and how is this relationship modulated by nitrogen fertilization? 3) How do microbial activity and the active microbiome vary between single-species and mixed-species treatments? 4) Which specific microbial taxa are enriched by each plant functional group, and does this enrichment persist in mixtures?

**Figure 1:**
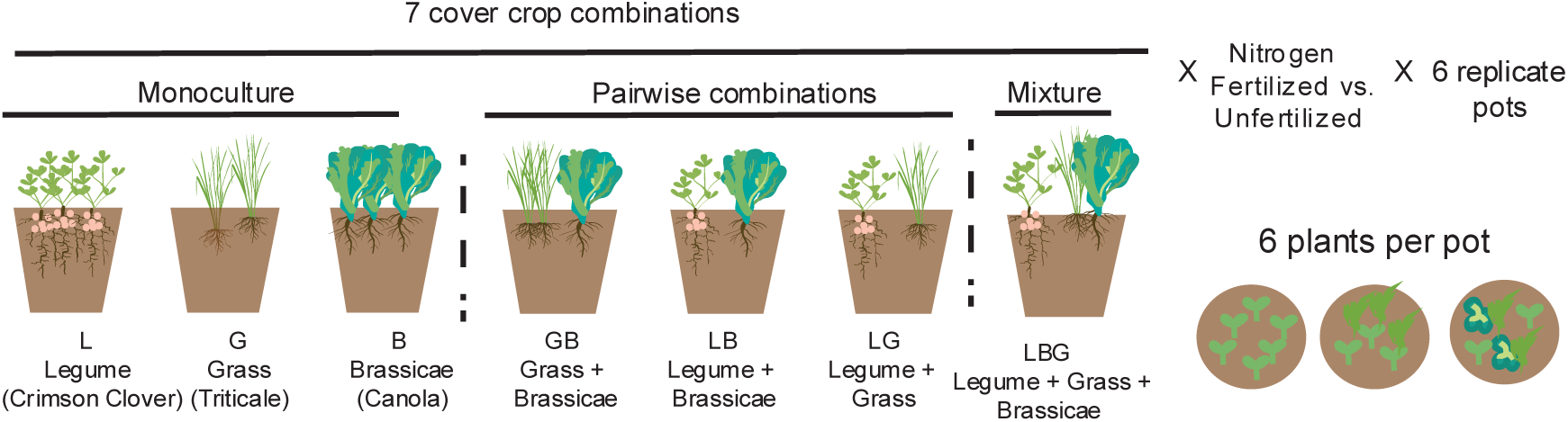
Schematic of experimental design, illustrating the seven cover crop combinations of monocultures, pairwise combinations, and three-species mixtures. Cover crop treatments consisted of two nitrogen levels and six replicates in a full factorial design. Replication was at the pot level with six plants per pot and 6 pots per treatment. Pots were randomized within each of the 6 blocks. Note that all the pots were harvested and processed to measure plant above and belowground biomass, nitrogen from fixation via isotopes, weed seed germination, and microbiome composition via rhizosphere Total 16S rRNA gene amplicon sequencing. BONCAT-FACS-Seq of rhizosphere active cell count and active and inactive microbiome community composition via 16S rRNA amplicon was performed only on the nitrogen-fertilized subset of plants.

## Material and methods

### Greenhouse experiment

We grew Crimson clover (legume, *Trifolium incarnatum*), canola (brassica, *Brassica napus*), and triticale (grass, *× Triticosecale*) in a factorial design with 7 plant treatments (monocultures, pairwise mixtures, three-species mixture) and 2 nitrogen levels (unfertilized vs. aqueous NH_4_NO_3_) with 6 replicates (N=84, Figure 1) randomized into four blocks, plus four unplanted soil control pots. Pots maintained a constant density of 6 plants per pot (6:0, 3:3, or 2:2:2). We refer to plant treatments by legume, brassica, grass, and their mixtures. Nitrogen-fertilized pots received 200 ppm aqueous NH_4_NO_3_ biweekly (300 ppm in week 12); unfertilized treatments only received water. Pots contained a 5:4:1 (v/v/v) mixture of field soil, sand (#1 Q-ROK, US Silica), and vermiculite. Field soil was sourced from the Russell E. Larson Agricultural Research Center (Rock Springs, PA, 40.7215, −77.9275), and had a previous history of a six-species cover crop mixture including all focal plants, details on the field rotation in Finney, White and Kaye, 2016, updates on seeding rates in Hosler *et al*., 2025. Baseline nutrient levels of soil in Table S1. Prior to sowing of the focal seeds, 400 foxtail (*Setaria faberi*) and 500 pigweed (*Amaranthus powellii*) seeds were mixed into each pot. Plants were grown for 12 weeks prior to harvesting and sample collection.

### BONCAT incubation and rhizosphere microbiome collection

We used BONCAT-FACS (Bioorthogonal Non-Canonical Amino Acid Tagging coupled to Fluorescence-Activated Cell Sorting) to quantify microbial activity and collect translationally active microbial cells. Following the methods of Harris *et al*., 2025, we dosed plants with aqueous HPG (L-Homopropargylglycine), a non-canonical methionine, in the greenhouse, and incubated them for 24-hours. All roots from each pot were gently shaken to discard loose soil and vortexed in 250 mL of sterile 1X phosphate-buffered saline (PBS) for 5 minutes to harvest rhizosphere cells. The rhizosphere suspension was cleared of roots and centrifuged at 30 x g for 3 minutes to pellet debris. The rhizosphere supernatant was aliquoted into 700 μL aliquots, mixed with 200 μL of 50% glycerol, and stored at -20 °C for BONCAT activity measurements and sorting. Additional rhizosphere aliquots were pelleted at 12000 g for 5 min and frozen for profiling of the total microbial community. Rhizosphere soil from the remaining suspensions was decanted after 48 hours and air-dried to constant weight. From aliquots, HPG-containing cells were pelleted, resuspended, and fluorescently labeled with the FAM picolyl azide dye (Click Chemistry Tools; Ex/Em 490/510) via copper-catalyzed click chemistry (Reichart *et al*. 2020). Active and inactive microbial cells from all seven plant treatments from nitrogen-fertilized pots and soil-only controls were quantified and sorted using a Beckman Coulter MoFlo Astrios EQ Cell Sorter. We quantified the percentage of active cells for 3 minutes per sample. Then, we collected all active microbial cells from a known volume and normalized them by grams of rhizosphere soil. See Supporting Information for details on BONCAT click chemistry and flow cytometry.

### Plant traits and seed germination

We measured total above- and belowground dry biomass. Nitrogen from biological nitrogen fixation was quantified in legume leaves from δ¹⁵N isotopic measurements (Blesh 2018; Peoples *et al*. 1995; Pinto *et al*. 2021). We measured foxtail and pigweed seed germination rates at harvest using standard petri plate assays. For biomass and seed germination, we calculated expected values in mixtures from measured traits in monocultures, analogous to calculating the relative yield total (De Witt 1960; Fowler 1982). All statistical analyses were conducted in R (version 4.5.1). We evaluated plant treatment effects on shoot and root biomass, percent nitrogen from fixation, and nitrogen fixed per legume in a linear model (aov function, stats package, R Core Team, 2025) with block as a fixed effect, and nitrogen level as an interaction effect. We conducted Tukey post hoc tests using estimated marginal means with the emmeans package (Russell V. Lenth and Julia Piaskowski 2026). We compared expected and measured shoot and root biomass for each mixture using a linear model with nitrogen level as an interaction effect and rep as a fixed effect. We confirmed normality using the Kolmogorov-Smirnov test and qq plots of the residuals. For weed seed germination, we used a similar approach, but with a binomial generalized linear model and a Tukey-Sidak post hoc test. Additional information on plant trait collection, expected value calculation, and statistical analysis is available in the Supporting Information.

### Microbiome sequencing and bioinformatic processing

We extracted DNA from the total rhizosphere community of all nitrogen-fertilized, unfertilized, and soil-only pots (86 Total DNA samples) using the DNeasy PowerSoil Pro Kit (Qiagen). We extracted DNA from the active and inactive fractions sorted using FACS (see BONCAT section above) of the rhizosphere microbiome from nitrogen-fertilized pots only (38 active and 37 inactive samples) by adapting the library prep protocol for cell fractions described by Reichart *et al*., 2020. Cells were lysed with prepGEM Bacteria Kit (New England Biolabs). We used 515f-Y (Parada 2016, 5′-GTGYCAGCMGCCGCGGTAA-3′) and 806R-B (Apprill 2015, 5’-GGACTACNVGGGTWTCTAAT-3’) primers to amplify the 16S rRNA gene. The Huck Life Science Genomics Core performed library preparation, indexing, sequencing, and demultiplexing on an Illumina NextSeq 2000 P1 (2 x 300 paired-end reads), yielding an average of 108,211 paired reads per sample.

Demultiplexed reads were processed using the QIIME 2 pipeline (v2022.2, Bolyen *et al*., 2019). Forward and reverse reads were trimmed to 220 bp, denoised, merged, and cleared of chimeras to generate amplicon sequence variants (ASVs) using DADA2 (Callahan *et al*. 2016) with expected errors set to 2 bp forward and 5 bp reverse, and the detection method to consensus. Taxonomy was assigned with a Naïve Bayes classifier pre-trained on Greengenes2 (McDonald *et al*. 2023). ASVs belonging to chloroplasts or mitochondria were removed with the R package Phyloseq, v1.44.0 (McMurdie & Holmes 2013). Due to divergent sampling processes and diversity levels, total rhizosphere and sorted datasets were separately checked for library saturation, and rarefied to 65,234 and 68,552 reads per sample, respectively, using Rrarefy (vegan package, v2.6-4, Oksanen, Jari *et al*., 2022).

### Microbiome statistical analysis

To assess treatment differences in the percentage of active microbial cells and the number of active cells (g^-1^ rhizosphere soil), we used binomial and negative binomial models, respectively, followed by Tukey-corrected post hoc tests. Shannon diversity and evenness metrics were calculated in Phyloseq (McMurdie & Holmes 2013), and treatment differences were assessed using an ANOVA followed by Tukey post hoc tests with nitrogen as an interaction effect. For beta diversity, we filtered out rare taxa with fewer than 5 mean reads across samples (<0.0001% abundance). Bray-Curtis dissimilarity matrices were analyzed using Canonical Analysis of Principal Coordinates (CAP) (Oksanen, Jari *et al*. 2022), using plant treatments and nitrogen as explanatory variables. The effect of the presence or absence of each plant functional group was assessed in a CAP (distance matrix ∼ legume * grass * brassica + block). We calculated expected microbiomes by multiplying each ASV’s relative abundance in a monoculture by the measured percentage of root biomass for that species in the mixture. We compared the expected and measured microbial composition by projecting the mixtures onto an index derived from the monocultures. To construct this index, we centered the log-transformed data, calculated the centroids of each monoculture, and then defined mixture points as scalar-vector projections along the vector from monoculture centroids. In pairwise mixtures, values are normalized by subtracting the expected value from the measured value for each replicate. To determine key microbial taxa that differ between plant communities, we selected the 20 ASVs with the highest species scores on CAP axis 1 and 2 in the CAP ordination of plant treatment and aggregated them to the lowest possible taxonomic level. We determined the differential abundance of these aggregated ASVs by the presence/absence of each plant functional group with DESeq2 (Love *et al*. 2014). Block was accounted for in all models; more details on microbiome statistical analysis can be found in the supplement.

## Results

### Plant trait responses to mixtures and monocultures

First, we evaluate the effect of plant treatment on shoot and root biomass, nitrogen from fixation, and weed seed germination. Shoot biomass of legumes is greater than that of any monoculture, with or without added nitrogen (Figure 2, Table S2-S4). In mixtures, shoot biomass is not different between measured and the additive expectation from monocultures, but it trends higher in the three-species mixture (Table S5; *F*_1,20_=3.90, *P*=0.063). Root biomass is greater in legumes than in any monoculture without nitrogen fertilizer, but when nitrogen is added, grass root biomass increases, resulting in root biomass similar between legumes and grass (Figure 2, Table S6-S8). For all legume mixtures, root biomass exceeds additive expectations from monocultures, with or without added nitrogen (Table S9). Next, the legume nitrogen from biological nitrogen fixation (BNF) increases from monoculture to mixtures with and without nitrogen fertilizer (Figure 2, Table S10-S12). When nitrogen is applied, the difference in nitrogen from BNF between monoculture and mixtures increases, values rise from monoculture to two-species mixtures and peak in the three-species mixture. Similarly, there is more nitrogen fixed per legume in the three-species mixtures than in monocultures, across both nitrogen fertilizer levels (Figure 2, Table S13-S15). Additionally, we explored seed germination for common weeds, foxtail (*Setaria faberi*) and pigweed (*Amaranthus powellii*). For foxtail, plant treatment has a small but statistically significant impact on seed germination, with lower germination in the three-species mixture than in any monoculture (Supplemental Figure S2, Table S16). For pigweed, we observe the lowest seed germination occurring in the brassica monoculture (Supplemental Figure S2, Table S17) and greater than expected seed germination in all brassica-containing mixtures (Table S19). In sum, while shoot biomass did not differ from expected values in mixtures, belowground outcomes such as root biomass, nitrogen fixation, and weed seed germination did.

**Figure 2:**
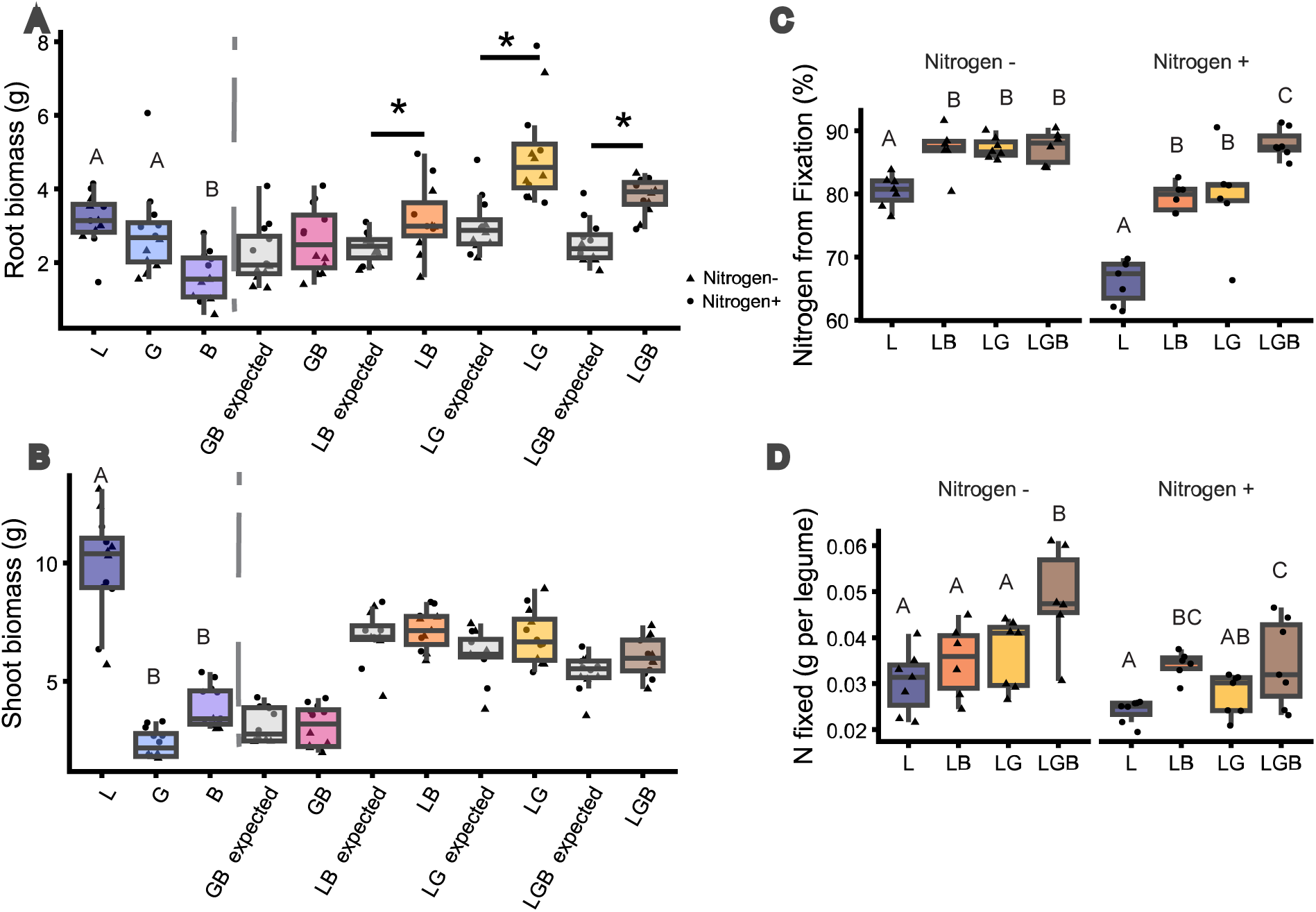
Plant traits for monocultures, additive expectations from monocultures, and mixtures. Measured and expected values for A) root biomass (g dry weight) and B) shoot biomass (g dry weight). Treatments labeled as expected (grey) are additive expectations from monocultures. Asterisks indicate significant differences between measured and expected values and from monoculture treatments in a one-way ANOVA. In plots A) and B), letters denote differences from a Tukey post hoc test that were significant both with and without added exogenous nitrogen. C) Percent nitrogen in legumes derived from biological nitrogen fixation versus chemical nitrogen sources. D) Grams of nitrogen fixed per legume plant. Both nitrogen levels are shown, with the shape indicating the nitrogen treatment. Both nitrogen levels are shown, with the shape indicating the nitrogen treatment. Letters denote differences from a Tukey post hoc test. Plant functional groups are abbreviated as L for legume, G for grass, and B for brassica. Mixtures are named using the abbreviations of all functional groups present; for example, LB indicates a legume-brassica mixture.

### Total rhizosphere microbiome responses to plant treatment and nitrogen fertilization

Next, we investigate how the plant treatment and nitrogen fertilization impact rhizosphere microbial composition. Overall, nitrogen fertilization decreases microbial Shannon diversity and ASV richness. Among monocultures, microbial Shannon diversity, ASV richness, and Pielou’s evenness are lower for legumes than for brassicas, but there are no differences between plant mixtures and monocultures (Figure S3, Tables S20-25). For beta diversity, there are significant effects of plant treatment and nitrogen fertilizer, and a marginal interaction between the two in Constrained Analysis of Principal Coordinates (CAP) of Bray Curtis dissimilarities (Figure S4; Table S26; *F*_6,68_ = 3.35, *P* = 0.001; *F*_1,68_ = 3.03, *P* = 0.001; and *F*_6,68_ = 1.18, *P* = 0.052). In both nitrogen-fertilized and unfertilized pots, we observe differences in beta diversity among legume, grass, and brassica monocultures (Figure 3; Tables S27-28). Without added nitrogen, legume presence has the largest effect on beta diversity (*F*_6,34_ = 6.49, *P* = 0.001), brassica does not have a significant effect, and brassica-containing mixtures are not different than grass or legume monocultures (Figure 3, Table S27-S29). In addition, we do not detect any significant interaction effects, indicating that effects are additive in mixtures without nitrogen fertilizer. When nitrogen is added, we find additional pairwise differences in beta diversity, with mixtures differing from their respective monocultures (Figure 3, Table S29). In nitrogen-fertilized pots, when we assess the contribution of each plant functional group to microbial composition in a CAP, we find a significant main effect of brassica and a significant legume × grass interaction, indicating non-additive effects when legumes and grasses are combined (Table S30). We explore whether differences in relative root biomass contribute to this non-additive effect by scaling values to monocultures and accounting for differences in the proportion of belowground biomass. Once root biomass is accounted for, we find that pairwise comparisons do not differ significantly from the additive expected values and are intermediate between their respective monocultures, although three-species mixtures are more similar to grass and less similar to brassica than expected (Figure 3, Table S31; PCA in Figure S5). Overall, the plant treatment minimally affects microbial alpha diversity but strongly influences microbial beta diversity in response to nitrogen addition, resulting in microbial communities in mixtures that are distinct from their respective monocultures, and often additive.

**Figure 3:**
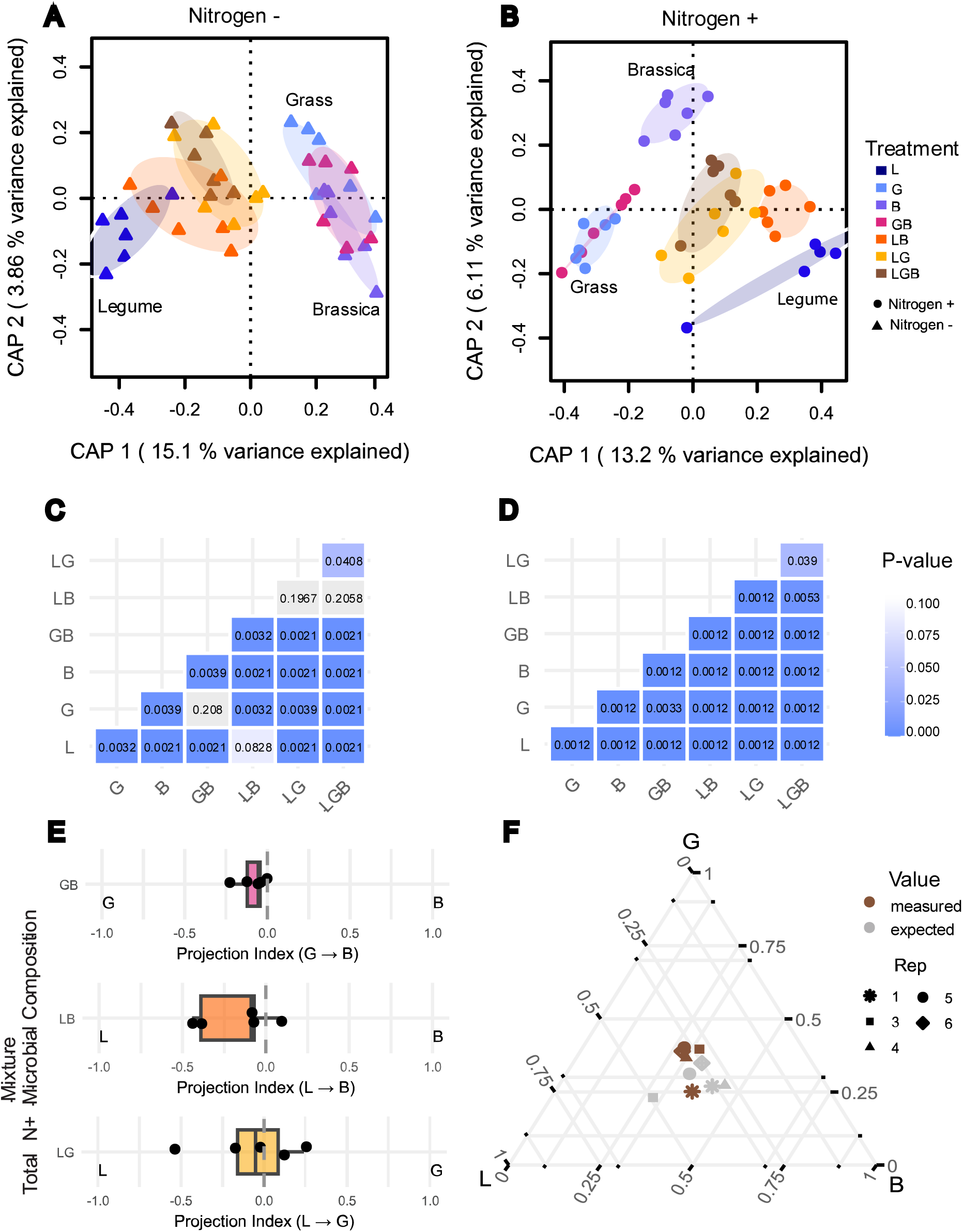
Rhizosphere total microbial composition in unfertilized (Nitrogen –) and fertilized (Nitrogen +) conditions. A-B) Beta diversity differences among plant treatments were tested and visualized with a Canonical Analysis of Principal Coordinates on Bray-Curtis dissimilarities in the total microbial community for samples A) without added nitrogen (n=6) and B) with added nitrogen (n=6). Ellipses are 95% confidence intervals. C-D) To determine which treatments differed in A and B, we performed a pairwise permutation test of the CAP model on Bray-Curtis dissimilarities (Table S27-28). Significant p-values are denoted by shades of blue. C) without added nitrogen, and D) with added nitrogen. E-F) To account for differences in relative root biomass between plant species, we projected the measured and expected microbial composition of mixtures for each replicate along an index between monocultures. To construct this index, we first center the log-transformed abundances, then define vectors between the centroids of each monoculture, and finally project mixture points as scalar-vector projections along those vectors (see Figure S5). E) To show differences from additive expectations in two-way mixtures, values are normalized so that zero is the expected value of each replicate, and the point is the difference between measured and expected. F) To show differences between measured and expected in the three-way mixtures, expected values are plotted in grey, measured values are brown, and the shape indicates replicate (Rep). The index for the 3-species mixture is not normalized; the raw values shown are between 0 and 1. For C-F, ASVs were rarefied, and extremely rare ASVs were removed. Plant functional groups are abbreviated as L for legume, G for grass, and B for brassica.

### Microbial activity and active microbial composition among monocultures and mixtures

Next, we assess non-additive effects of plant composition on the active microbiome by measuring microbial activity and active microbial composition in the rhizosphere under nitrogen fertilization (Figure 4). We measure microbial activity by the number of BONCAT-active cells per gram of rhizosphere and by the percentage of active cells among total cells. Across both indicators, legume-containing treatments show increased microbial activity (Tables S32-33; z = 1.59, SE = 0.25, *P* < 0.001 and z = 0.06, SE =0.01, *P* < 0.001). Active cells per gram of rhizosphere increase in legumes compared with other monocultures, and the expected and measured values for the mixtures were not statistically different (Figure 4, Table S34-35). For the percentage of active cells, the observed value exceeds that expected from monocultures in the LG pairwise mixtures (Table S36-37; z = 2.55, SE = 0.23, *P* = 0.010). Overall, the presence of legumes increases microbial activity, with non-additive effects in the percentage of active cells in legume-grass mixtures.

**Figure 4:**
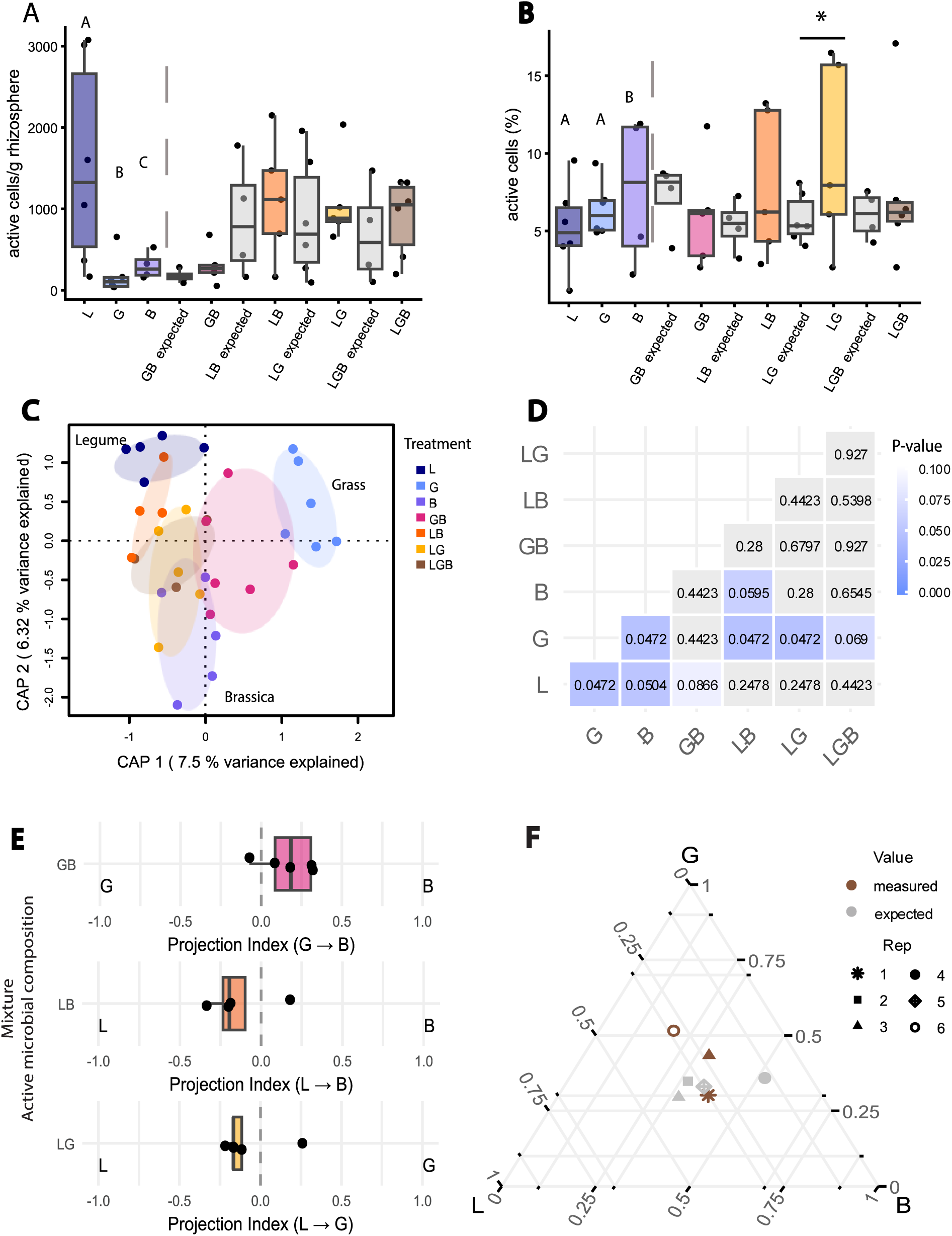
Microbial activity and active microbial community composition responses to plant treatment. Active microbiomes were sequenced only from nitrogen-fertilized treatments due to low bacterial cell counts. A) Treatment effects on active microbial cell count per gram of rhizosphere soil. Letters indicating differences in a Tukey post hoc test using a negative binomial model are shown for the monocultures (Table S33). Mixtures did not differ significantly from expectations based on monocultures (Table S35). Soil-only treatment is not shown (mean 827 ± SD 296). B) Treatment effects on the percent active microbial cells. Letters indicating differences in a Tukey post hoc test using a binomial model are shown for the monocultures (Table S34). For mixtures, asterisks indicate significant differences in a one-way ANOVA between measured and expected values. Soil-only treatment is not shown (mean 1.7% ± SD 1.3). C) Beta diversity differences among plant treatments are tested and visualized with a Canonical Analysis of Principal Coordinates (CAP) on Bray-Curtis dissimilarities in the active community. Ellipses denote 95% confidence intervals. D) To determine which treatments differ in C, we perform a pairwise permutation test of the CAP model on Bray-Curtis dissimilarities (Table S38). Significant p-values are denoted by shades of blue. E-F) To account for differences in relative root biomass between plant species, we projected the measured and expected microbial composition of mixtures for each replicate along an index between monocultures. To construct this index, we first center the log-transformed abundances, then define vectors between the centroids of each monoculture, and finally project mixture points as scalar-vector projections along those vectors (see Figure S5). E) To show differences from additive expectations in two-way mixtures, values are normalized so that zero is the expected value of each replicate, and the point is the difference between measured and expected. F) To show differences between measured and expected in the three-way mixtures, expected values are plotted in grey, measured values are brown, and the shape indicates replicate. The index for the 3-species mixture is not normalized; the raw values shown range from 0 to 1. For C-F, ASVs were rarefied, and extremely rare ASVs were removed. Plant functional groups are abbreviated as L for legume, G for grass, and B for brassica.

Then, we evaluate how the active microbial composition varies among plant treatments. The active and inactive microbial community structure differ in the rhizosphere, as shown by the CAP of Bray-Curtis dissimilarities (Figure 4C, Table S38; *F*_1,52_ = 4.28, *P* = 0.001). In both active and inactive communities, plant treatment does not significantly affect microbial alpha diversity (Supplemental Figure S6), but plant treatment does significantly affect beta diversity for both active (Figure 4D, Table S38; *F*_6,52_ = 1.77, *P* = 0.001) and inactive communities (Supplemental Figure S7 and Table S38-39). In the active community, plant monocultures are distinct (Table S40), and the grass monoculture is distinct from the legume-grass mixture, and the brassicae monoculture is distinct from the legume-brassica mixture (Table S40). Overall, the plant treatment affects the active microbiome, but more subtly than in the total microbiome.

In the active microbial community, we explore the extent to which community composition in mixtures differs from additive expectations. In a CAP of the effect of each plant functional group on the active community beta diversity, there is a significant legume × grass interaction (Table S41), indicating non-additive effects when legumes and grass are combined. When scaled relative to monocultures and after accounting for differences in root biomass between species, mixtures fall between their respective monocultures and do not differ from the expected values (Figure 4, panels E and F; PCA in Figure S8; Table S42). In sum, like the total microbiome, when differences in belowground biomass are accounted for, the active microbiome composition is largely additive.

### Differential abundance analysis of specific taxa in response to plant treatments

Because there are significant main and interaction effects of plant treatments on beta diversity in both total and active samples, we assess the individual taxa that contribute most strongly to differences in microbiome composition in a differential abundance analysis with DESeq2. First, we examine focal taxa in the total microbiome. We find multiple ASVs of the genus *Rhizobium,* which symbiotically fix nitrogen with the legume species in the experiment, increase when legumes are present, but show significant legume × brassica and legume × grass interactions (Figure 5, Table S43). We also find that ASVs of the ammonia-oxidizing archaeon *Nitrosocosmicus spp.* have the highest abundance when brassica is present, with significant grass × brassica interaction and values decreasing when brassica and grass were combined (Table S43), and that *Caulobacter spp.,* many of which are plant growth-promoting bacteria, increase in all plant treatments, with the largest increase in legume treatments (Table S43). Lastly, the presence of grass increases the abundance of *Polaromonas sp.*, a suspected associative nitrogen fixer (Table S43). Second, we examine focal taxa in the active community. We find several ASVs from the phylum Actinomycetes, a diverse phylum common in soils, that decrease when brassica is present and whose abundance is altered when legumes and grasses interact (Table S44). In addition, we found that ASVs in the genus *Vampirovibrio* (phylum Cyanobacterium), which primarily consist of predatory bacteria, show a significant legume × grass interaction (Table S44). The abundance of *Deinococcus sp*., a bacterium highly resistant to harsh environmental conditions, increases in the presence of brassica (Table S44). We also find that taxa with significant patterns in the total community were present in the active community. Overall, in the total microbiome, we find many microbial taxa whose abundance was significantly affected by plant species combinations, whereas in the active subset, we find a handful of taxa influenced by brassica and/or legume-grass combinations.

**Figure 5.**
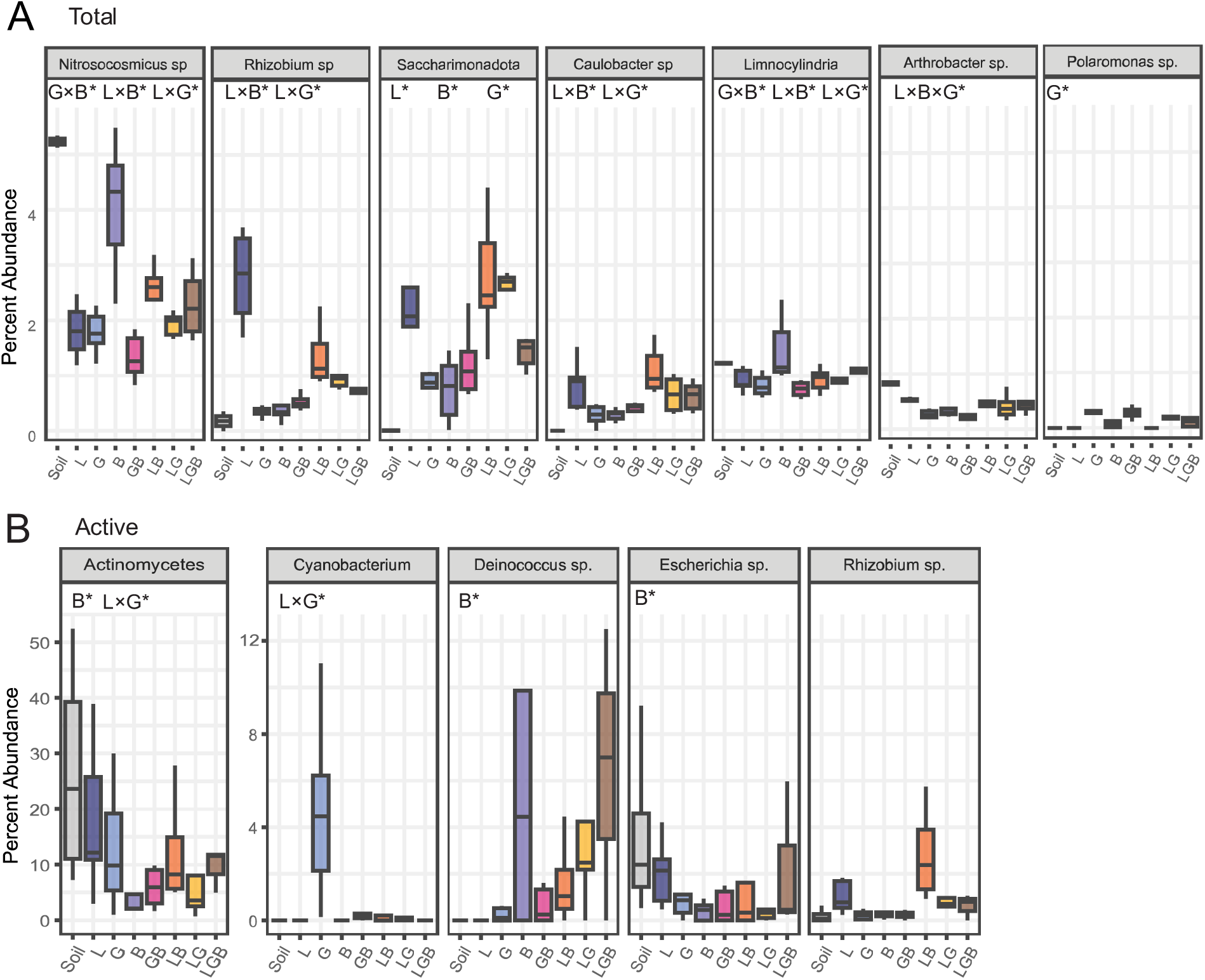
Differential abundance among plant functional groups of key taxa from the Total (A) and Active subset (B) of the rhizosphere microbial community. First, the twenty ASVs with the highest species scores on CAP 1 and CAP 2 from our CAP ordination were selected from the active and total communities. These ASVs were aggregated to the lowest taxonomic level, typically the genus level. Then DESeq2 was run on these key taxa (formula: abundance ∼ Legume * Grass *Brassica). Model terms are the binary variables of the present/absence of a plant functional group in a plant mixture. Letters on graphs indicate significant effects for each taxon in a DESeq2 model (full results in Tables S41 and S42). Taxa without letters did not differ significantly among plant functional groups in DESeq2.

## Discussion

We evaluated whether plant functional group mixtures are additive intermediates of their monocultures across plant traits, total and active microbiomes. In monoculture, plant functional groups had distinct characteristics: legume monocultures had the greatest shoot and root biomass and the highest abundance of active microbial cells; brassica most effectively suppressed pigweed germination; and all three monocultures had distinct rhizosphere microbial communities. In mixtures, non-additive effects emerged, including higher-than-expected root biomass, nitrogen from biological nitrogen fixation (BNF) in legume mixtures, and higher microbial activity in legume-grass combinations. Furthermore, under nitrogen fertilization, total and active microbial communities exhibited non-additive effects when legumes and grasses were combined, affecting specific taxa such as *Rhizobium sp*. and *Caulobacter sp.* These non-additive effects in the microbiome were facilitated by differences in the proportion of belowground biomass. When adjusted for the proportion of belowground biomass and scaled between monocultures, communities in mixtures fall between their respective monocultures. Overall, while legume mixtures can non-additively increase nitrogen fixation, root biomass, and microbial activity, microbial community assembly in mixtures is predictable from the contributions of belowground plant biomass.

### Plant belowground biomass and nitrogen fixation are higher in legume mixtures than expected from monocultures

Biomass was highest in legumes and their mixtures, likely due to biological nitrogen fixation (BNF). Soils were likely nitrogen-limited because there was substantial BNF in all legume samples, even in nitrogen-fertilized treatments. In these conditions, legumes could overcome nitrogen limitation by relying on symbiotic nitrogen-fixing rhizobia and produce greater biomass than non-legumes, as BNF and biomass are associated (Blesh 2018). In legume mixtures, effective nitrogen scavengers (Dean & Weil 2009; Hirsh *et al*. 2021; Kaye *et al*. 2019), outcompeted legumes for soil nitrogen, forcing legumes to increase BNF. Increased nitrogen in the system likely contributed to the non-additive increases in root biomass in legume mixtures. Our results align with other studies that show non-additive increases in legume nitrogen pools in mixtures (Roscher *et al*. 2008; Spehn *et al*. 2002). Although the magnitude of BNF decreased under nitrogen addition (Reinprecht *et al*. 2020; Salvagiotti *et al*. 2009; Waterer & Vessey 1993), our results indicate that the mixtures can stimulate additional nitrogen fixed relative to monocultures, increasing belowground productivity in the legume mixtures.

### Nitrogen availability regulates the effects of plant composition on the microbiome

We found that plant composition strongly shifts microbiome structure, especially under nitrogen fertilization, while having a limited effect on microbial alpha diversity. Although we expected alpha diversity to increase with plant richness (Schmid *et al*. 2019), mixtures may not have been sufficiently diverse, or the experiment may have been too brief to create detectable changes. In contrast, beta diversity diverged among plant monocultures, corroborating established patterns in the rhizosphere (Haichar *et al*. 2008; Lay *et al*. 2018) and bulk soil (Zhou *et al*. 2017). Beta diversity also differed between monocultures and mixtures. While such differences are inconsistently observed in bulk soil (Maciá-Vicente *et al*. 2024; Ryan *et al*. 2026; Schmid *et al*. 2019), sampling the rhizosphere likely increased our sensitivity to these shifts (Lopes *et al*. 2021). Furthermore, our results alongside recent work (Isbell *et al*. 2026) suggest that variable outcomes may stem from an interaction between nitrogen and plant composition on the microbiome. Plants need nitrogen to produce costly root exudates that shape microbiomes (Wen *et al*. 2022; Yang *et al*. 2025; Zhu *et al*. 2016). Only with nitrogen addition do we find that brassica alters microbial composition, and non-additive legume × grass effects emerge. Our results show that sufficient nitrogen is required for non-legume plant species to significantly shift microbiome composition.

### Non-additive shifts in legume grass mixtures are mediated by belowground biomass

Under nitrogen enrichment, the total and active rhizosphere microbiomes exhibited similar patterns, characterized by functional-group divergence and non-additive responses in legume grass mixtures. However, the metabolically active fraction revealed fewer significant differences than the total community. This difference may be due to only measuring active microbes during a 24-hour incubation, and microbial composition can change substantially during plant development (Houlden *et al*. 2008). In both the total and active microbiome, brassicas had the weakest effect on microbial composition, despite brassicas’ ability to produce allelopathic compounds and glucosinolates (Haramoto & Gallandt 2004; Petersen *et al*. 2001). Instead, legumes had the greatest impact on microbial composition, in line with Zhou *et al*. 2017, and legumes increased microbial activity. In mixtures, microbial activity and composition were not purely additive. Legume-grass mixtures had higher-than-expected microbial activity, and across both the active and total microbiomes, they exhibited non-additive shifts in composition, potentially contributing to the increased plant productivity observed in mixtures (Schnitzer et al., 2011). However, when we adjust for differences in belowground plant biomass, we find that mixture compositions are intermediate between monocultures, suggesting that non-additive effects are, in part, attributable to these belowground biomass differences. Although other studies have not predicted microbial composition with plant biomass, plant biomass predicts microbial biomass reasonably well (Fierer *et al*. 2009). Our results suggest that incorporating plant belowground biomass can account for non-additive effects in microbial composition in mixtures, which could improve the predictability of microbiomes.

### Plant functional groups enrich distinct microbial taxa involved in plant growth promotion and the nitrogen cycle

Understanding the microbes associated with each plant functional group, both individually and in mixtures, also offers opportunities to manage microbial-mediated functions. *Caulobacter sp.,* a common plant-associated genus with some strains capable of promoting plant growth (Berrios & Ely 2020; Edulamudi *et al*. 2011), increased in all plant treatments, with the largest increase occurring in legume mixtures. Next, we find that brassica in monocultures increases the abundance of *Nitrosocosmicus sp.*, a clade of ammonia-oxidizing archaea often involved in nitrogen cycling across multiple environments (Han *et al*. 2024). Interestingly, this positive effect of brassica is nullified in grass brassica mixtures, indicated by a significant grass × brassica interaction. This suggests that growing brassicas could increase ammonia oxidation, but not in grass mixtures. In legume monocultures and mixtures, we observe an increase in the abundance of *Rhizobium* sp., consistent with other studies indicating that legumes can increase the abundance of their nitrogen-fixing symbionts in the soil (Chemining’wa & Vessey 2006; Kucey & Hynes 1989). In plant mixtures with grass, we find small but significant increases in the abundance of *Polaromonas* sp. *Polaromonas sp*. shows potential as an associative nitrogen fixer, as they harbor nitrogen-fixing genes in the literature, and increase in grass treatments (Darcy *et al*. 2011; Hanson *et al*. 2012; Nash *et al*. 2018; Sun *et al*. 2010). Growing each of these plant species, in monoculture or in combination, could promote these potentially beneficial organisms.

### Conclusions and future directions

Although many microbial responses in mixtures were additive, certain combinations, such as legume-grass mixtures, generated non-additive shifts in active and total microbial communities that coincided with enhanced root biomass and nitrogen fixation. At the whole-community level, we find that non-additive effects in the microbiome were largely due to disproportionate belowground biomass, and that, when we account for it, mixtures can serve as predictive intermediates of monocultures. By identifying plant combinations that consistently promote beneficial microbial activity, it may be possible to design cover crop mixtures and restoration plantings that enhance nutrient cycling and plant productivity through targeted manipulation of the rhizosphere microbiome.

## Acknowledgments

The co-authors acknowledge the Huck Institutes’ Flow Cytometry Core Facility (RRID: SCR_024460) for use of the BD Fortessa Flow Cytometer and staff Mitchell Koptchak for collecting the data shown in Figure 4, and lab members for their helpful feedback. This work is supported by the USDA National Institute of Food and Agriculture and Hatch Appropriations under Project #PEN04949 and Accession #7006508 to E.M.C. and Project #PEN04760 and Accession #1025611 to L.T.B. This work is supported NIFA Predoctoral Fellowship, Award #: 2023-67011-40517 to J.E.H. and Award #: 2023-67011-40329 to E.K.R from the U.S. Department of Agriculture’s National Institute of Food and Agriculture and a seed grant from the Penn State College of Agricultural Sciences.

## Author Contributions

JEH was responsible for formal analysis, visualization, investigation, and writing of the original draft, with input from all authors. JEH and ER oversaw the project execution, methodology, and data curation. ER acquired data on nitrogen fixation, plant biomass, and seed germination, and JEH acquired data on microbial composition and activity. CL, EC, and LTB contributed to the conceptualization, funding, supervision, and execution of the project. All authors reviewed the manuscript.

## Data Availability Statement

The raw sequencing reads are available at the National Center for Biotechnology Information (NCBI) Sequence Read Archive under SRA16432418 and BioProject PRJNA1518164. Data used for analysis is available at the USDA National Agriculture Library Ag Data Commons https://doi.org/10.15482/USDA.ADC/33324813. The R code for analysis is available at https://github.com/jennnnnharris/.

## References

Aires, A., Mota, V.R., Saavedra, M.J., Monteiro, A.A., Simões, M., Rosa, E.A.S., et al. (2009). Initial *in vitro* evaluations of the antibacterial activities of glucosinolate enzymatic hydrolysis products against plant pathogenic bacteria. J. Appl. Microbiol., 106, 2096–2105.

Batstone, R.T., Ibrahim, A. & MacLean, L.T. (2023). Microbiomes: Getting to the root of the rhizobial competition problem in agriculture. Curr. Biol., 33, R777–R780.

Berrios, L. & Ely, B. (2020). Plant growth enhancement is not a conserved feature in the Caulobacter genus. Plant Soil, 449, 81–95.

Bertness, M.D. & Callaway, R. (1994). Positive interactions in communities. Trends Ecol. Evol., 9, 191– 193.

Bever, J.D., Westover, K.M. & Antonovics, J. (1997). Incorporating the Soil Community into Plant Population Dynamics: The Utility of the Feedback Approach. J. Ecol., 85, 561.

Blagodatskaya, E. & Kuzyakov, Y. (2013). Active microorganisms in soil: Critical review of estimation criteria and approaches. Soil Biol. Biochem., 67, 192–211.

Blanco-Canqui, H. (2024). Do cover crop mixtures improve soil physical health more than monocultures? Plant Soil, 495, 99–112.

Blesh, J. (2018). Functional traits in cover crop mixtures: Biological nitrogen fixation and multifunctionality. J. Appl. Ecol., 55, 38–48.

Bolyen, E., Rideout, J.R., Dillon, M.R., Bokulich, N.A., Abnet, C.C., Al-Ghalith, G.A., et al. (2019). Reproducible, interactive, scalable and extensible microbiome data science using QIIME 2. Nat. Biotechnol., 37, 852–857.

Bowman, M., Ferraro, P.J., Fuller, K.B., Gramig, B.M., Mosheim, R., Njuki, E., et al. (2025). Economic outcomes of soil health and conservation practices on U.S. cropland. U.S. Department of Agriculture, Economic Research Service, [Washington, D.C.].

Callahan, B.J., McMurdie, P.J., Rosen, M.J., Han, A.W., Johnson, A.J.A. & Holmes, S.P. (2016). DADA2: High-resolution sample inference from Illumina amplicon data. Nat. Methods, 13, 581–583.

Carini, P., Marsden, P.J., Leff, J.W., Morgan, E.E., Strickland, M.S. & Fierer, N. (2016). Relic DNA is Abundant in Soil and Obscures Estimates of Soil Microbial Diversity. Nat. Microbiol., 2, 16242.

Chemining’wa, G.N. & Vessey, J.K. (2006). The abundance and efficacy of Rhizobium leguminosarum bv. viciae in cultivated soils of the eastern Canadian prairie. Soil Biol. Biochem., 38, 294–302.

Cloutier, M.L., Murrell, E., Barbercheck, M., Kaye, J., Finney, D., García-González, I., et al. (2020). Fungal community shifts in soils with varied cover crop treatments and edaphic properties. Sci. Rep., 10, 6198.

Couradeau, E., Sasse, J., Goudeau, D., Nath, N., Hazen, T.C., Bowen, B.P., et al. (2019). Probing the active fraction of soil microbiomes using BONCAT-FACS. Nat. Commun., 10, 2770.

Darcy, J.L., Lynch, R.C., King, A.J., Robeson, M.S. & Schmidt, S.K. (2011). Global Distribution of Polaromonas Phylotypes - Evidence for a Highly Successful Dispersal Capacity. PLoS ONE, 6, e23742.

De Witt, C.T. (1960). On competition. Wageningen university.

Dean, J.E. & Weil, R.R. (2009). Brassica Cover Crops for Nitrogen Retention in the Mid-Atlantic Coastal Plain. J. Environ. Qual., 38, 520–528.

Dee, L.E., Ferraro, P.J., Severen, C.N., Kimmel, K.A., Borer, E.T., Byrnes, J.E.K., et al. (2023). Clarifying the effect of biodiversity on productivity in natural ecosystems with longitudinal data and methods for causal inference. Nat. Commun., 14, 2607.

Edulamudi, P., Antony Masilamani, A.J., Divi, V.R.S.G. & Konada, V.M. (2011). Novel root nodule bacteria belonging to the genus Caulobacter: Novel root nodule bacteria belonging to Caulobacter. Lett. Appl. Microbiol., 53, 587–591.

Fierer, N., Strickland, M.S., Liptzin, D., Bradford, M.A. & Cleveland, C.C. (2009). Global patterns in belowground communities. Ecol. Lett., 12, 1238–1249.

Finney, D.M., Buyer, J.S. & Kaye, J.P. (2017). Living cover crops have immediate impacts on soil microbial community structure and function. J. Soil Water Conserv., 72, 361–373.

Finney, D.M. & Kaye, J.P. (2017). Functional diversity in cover crop polycultures increases multifunctionality of an agricultural system. J. Appl. Ecol., 54, 509–517.

Finney, D.M., White, C.M. & Kaye, J.P. (2016). Biomass Production and Carbon/Nitrogen Ratio Influence Ecosystem Services from Cover Crop Mixtures. Agron. J., 108, 39–52.

Florence, A.M. & McGuire, A.M. (2020). Do diverse cover crop mixtures perform better than monocultures? A systematic review. Agron. J., 112, 3513–3534.

Fowler, N. (1982). Competition and Coexistence in a North Carolina Grassland: III. Mixtures of Component Species. J. Ecol., 70, 77.

Groff, S. (2015). The past, present, and future of the cover crop industry. J. Soil Water Conserv., 70, 2.

Haichar, F.E.Z., Marol, C., Berge, O., Rangel-Castro, J.I., Prosser, J.I., Balesdent, J., et al. (2008). Plant host habitat and root exudates shape soil bacterial community structure. ISME J., 2, 1221–1230.

Han, S., Kim, S., Sedlacek, C.J., Farooq, A., Song, C., Lee, S., et al. (2024). Adaptive traits of *Nitrosocosmicus* clade ammonia-oxidizing archaea. mBio, 15, e02169–24.

Hanson, B.T., Yagi, J.M., Jeon, C.O. & Madsen, E.M. (2012). Role of nitrogen fixation in the autecology of *Polaromonas naphthalenivorans* in contaminated sediments. Environ. Microbiol., 14, 1544– 1557.

Haramoto, E.R. & Gallandt, E.R. (2004). Brassica cover cropping for weed management: A review. Renew. Agric. Food Syst., 19, 187–198.

Harris, J.E., Bledsoe, R.B., Guha, S., Omari, H., Crandall, S.G., Burghardt, L.T., et al. (2025). The activity of soil microbial taxa in the rhizosphere predicts the success of root colonization. mSystems, 10, e00458–25.

Hector, A., Schmid, B., Beierkuhnlein, C., Caldeira, M.C., Diemer, M., Dimitrakopoulos, P.G., et al. (1999). Plant Diversity and Productivity Experiments in European Grasslands. Science, 286, 1123–1127.

Hendriks, M., Mommer, L., De Caluwe, H., Smit-Tiekstra, A.E., Van Der Putten, W.H. & De Kroon, H. (2013). Independent variations of plant and soil mixtures reveal soil feedback effects on plant community overyielding. J. Ecol., 101, 287–297.

Hirsh, S.M., Duiker, S.W., Graybill, J., Nichols, K. & Weil, R.R. (2021). Scavenging and recycling deep soil nitrogen using cover crops on mid-Atlantic, USA farms. Agric. Ecosyst. Environ., 309, 107274.

Hooper, D.U., Chapin, F.S., Ewel, J.J., Hector, A., Inchausti, P., Lavorel, S., et al. (2005). Effects of Biodiversity on Ecosystem Functioning: A Consensus of Current Knowledge. Ecol. Monogr., 75, 3–35.

Hooper, D.U. & Dukes, J.S. (2004). Overyielding among plant functional groups in a long-term experiment. Ecol. Lett., 7, 95–105.

Hosler, S.C., Murrell, E.G., Arrington, K.E., Baraibar, B., Barbercheck, M.E., Bradley, B.A., et al. (2025). Managing cover crop mixtures over a decade via species replacement and seeding rate adjustment. Agric. Environ. Lett., 10, e70029.

Houlden, A., Timms-Wilson, T.M., Day, M.J. & Bailey, M.J. (2008). Influence of plant developmental stage on microbial community structure and activity in the rhizosphere of three field crops: Plant and growth stage effects on microbial populations. FEMS Microbiol. Ecol., 65, 193–201.

Huston, M.A. (1997). Hidden treatments in ecological experiments: re-evaluating the ecosystem function of biodiversity. Oecologia, 110, 449–460.

Isbell, S.A., Reardon, K.M., Yates, C., Bell, T.H. & Kaye, J.P. (2026). Cover Crop Influence over Soil Microbiomes Varies by Plant Functional Group and Is Nutrient Dependent. Phytobiomes J., PBIOMES-08-25-0062-SC.

Jiang, P., Wang, Y., Zhang, Y., Fei, J., Rong, X., Peng, J., et al. (2024). Intercropping enhances maize growth and nutrient uptake by driving the link between rhizosphere metabolites and microbiomes. New Phytol., 243, 1506–1521.

Kaye, J., Finney, D., White, C., Bradley, B., Schipanski, M., Alonso-Ayuso, M., et al. (2019). Managing nitrogen through cover crop species selection in the U.S. mid-Atlantic. PLOS ONE, 14, e0215448.

Kou, T.J., Zhu, P., Huang, S., Peng, X.X., Song, Z.W., Deng, A.X., et al. (2012). Effects of long-term cropping regimes on soil carbon sequestration and aggregate composition in rainfed farmland of Northeast China. Soil Tillage Res., 118, 132–138.

Kucey, R.M.N. & Hynes, M.F. (1989). Populations of *Rhizobium leguminosarum* biovars *phaseoli* and *viceae* in fields after bean or pea in rotation with nonlegumes. Can. J. Microbiol., 35, 661–667.

Lay, C.-Y., Bell, T.H., Hamel, C., Harker, K.N., Mohr, R., Greer, C.W., et al. (2018). Canola Root– Associated Microbiomes in the Canadian Prairies. Front. Microbiol., 9, 1188.

Lopes, L.D., Hao, J. & Schachtman, D.P. (2021). Alkaline soil pH affects bulk soil, rhizosphere and root endosphere microbiomes of plants growing in a Sandhills ecosystem. FEMS Microbiol. Ecol., 97, fiab028.

Love, M.I., Huber, W. & Anders, S. (2014). Moderated estimation of fold change and dispersion for RNA-seq data with DESeq2. Genome Biol., 15, 550.

Maciá-Vicente, J.G., Cazzaniga, S., Duhamel, M., Van Den Beld, L., Lombaers, C., Visser, J., et al. (2024). Cover crop mixtures do not assemble markedly distinct soil microbiotas as compared to monocultures in a multilocation field experiment. Appl. Soil Ecol., 202, 105573.

McDonald, D., Jiang, Y., Balaban, M., Cantrell, K., Zhu, Q., Gonzalez, A., et al. (2023). Greengenes2 unifies microbial data in a single reference tree. Nat. Biotechnol.

McMurdie, P.J. & Holmes, S. (2013). phyloseq: An R Package for Reproducible Interactive Analysis and Graphics of Microbiome Census Data. PLoS ONE, 8, e61217.

Mueller, K.E., Tilman, D., Fornara, D.A. & Hobbie, S.E. (2013). Root depth distribution and the diversity–productivity relationship in a long-term grassland experiment. Ecology, 94, 787–793.

Nair, A. & Ngouajio, M. (2012). Soil microbial biomass, functional microbial diversity, and nematode community structure as affected by cover crops and compost in an organic vegetable production system. Appl. Soil Ecol., 58, 45–55.

Nash, M.V., Anesio, A.M., Barker, G., Tranter, M., Varliero, G., Eloe-Fadrosh, E.A., et al. (2018). Metagenomic insights into diazotrophic communities across Arctic glacier forefields. FEMS Microbiol. Ecol., 94.

Oksanen, Jari, Simpson, Gavin L., Blanchet, F. Guillaume, Kindt, Roeland, Legendre, Pierre, Minchin, Peter R., et al. (2022). Vegan: Community Ecology Package. R Package Version 26-4.

Peoples, M.B., Herridge, D.F. & Ladha, J.K. (1995). Biological nitrogen fixation: An efficient source of nitrogen for sustainable agricultural production? Plant Soil, 174, 3–28.

Petersen, J., Belz, R., Walker, F. & Hurle, K. (2001). Weed Suppression by Release of Isothiocyanates from Turnip-Rape Mulch. Agron. J., 93, 37–43.

Pinto, P., Rubio, G., Gutiérrez, F., Sawchik, J., Arana, S. & Piñeiro, G. (2021). Variable root:shoot ratios and plant nitrogen concentrations discourage using just aboveground biomass to select legume service crops. Plant Soil, 463, 347–358.

R Core Team. (2025). R: A Language and Environment for Statistical Computing.

Reichart, N.J., Jay, Z.J., Krukenberg, V., Parker, A.E., Spietz, R.L. & Hatzenpichler, R. (2020). Activity-based Cell Sorting Reveals Responses of Uncultured Archaea and Bacteria to Substrate Amendment. ISME J., 14, 2851–2861.

Reinprecht, Y., Schram, L., Marsolais, F., Smith, T.H., Hill, B. & Pauls, K.P. (2020). Effects of Nitrogen Application on Nitrogen Fixation in Common Bean Production. Front. Plant Sci., 11, 1172.

Richards, S.C., King, W.L., Cao, L.Y., Bradley, B.A., Rice, E.K., Lowry, C.J., et al. (2026). Microbial Colonizers in an Agroecosystem Under Diverse Cover Crop Treatments. Phytobiomes J., 10, 83– 97.

Rodríguez, H., Fraga, R., Gonzalez, T. & Bashan, Y. (2006). Genetics of phosphate solubilization and its potential applications for improving plant growth-promoting bacteria. Plant Soil, 287, 15–21.

Roscher, C., Thein, S., Schmid, B. & Scherer-Lorenzen, M. (2008). Complementary nitrogen use among potentially dominant species in a biodiversity experiment varies between two years. J. Ecol., 96, 477–488.

Russell V. Lenth and Julia Piaskowski. (2026). emmeans: Estimated Marginal Means, aka Least-Squares Means.

Ryan, K.B., Finn, J.A., De Menezes, A., Byrne, L., Brophy, C. & Brennan, F.P. (2026). Plant Identity Impacts the Soil Microbiome More Than Interspecific Interactions in Intensively Managed Grasslands. Eur. J. Soil Sci., 77, e70256.

Salvagiotti, F., Specht, J.E., Cassman, K.G., Walters, D.T., Weiss, A. & Dobermann, A. (2009). Growth and Nitrogen Fixation in High-Yielding Soybean: Impact of Nitrogen Fertilization. Agron. J., 101, 958–970.

Schimel, J.P. & Bennett, J. (2004). Nitrogen Mineralization: Challenges of a Changing Paradigm. Ecology, 85, 591–602.

Schmid, M.W., Hahl, T., Van Moorsel, S.J., Wagg, C., De Deyn, G.B. & Schmid, B. (2019). Feedbacks of plant identity and diversity on the diversity and community composition of rhizosphere microbiomes from a long-term biodiversity experiment. Mol. Ecol., 28, 863–878.

Schnitzer, S.A., Klironomos, J.N., HilleRisLambers, J., Kinkel, L.L., Reich, P.B., Xiao, K., et al. (2011). Soil microbes drive the classic plant diversity–productivity pattern. Ecology, 92, 296–303.

Singh, A., Ghimire, R. & Acharya, P. (2024). Soil profile carbon sequestration and nutrient responses varied with cover crops in irrigated forage rotations. Soil Tillage Res., 238, 106020.

Snapp, S.S., Swinton, S.M., Labarta, R., Mutch, D., Black, J.R., Leep, R., et al. (2005). Evaluating Cover Crops for Benefits, Costs and Performance within Cropping System Niches. Agron. J., 97, 322– 332.

Spehn, E.M., Scherer-Lorenzen, M., Schmid, B., Hector, A., Caldeira, M.C., Dimitrakopoulos, P.G., et al. (2002). The role of legumes as a component of biodiversity in a cross-European study of grassland biomass nitrogen. Oikos, 98, 205–218.

Sun, W., Xie, S., Luo, C. & Cupples, A.M. (2010). Direct Link between Toluene Degradation in Contaminated-Site Microcosms and a *Polaromonas* Strain. Appl. Environ. Microbiol., 76, 956– 959.

Tilman, D., Knops, J., Wedin, D., Reich, P., Ritchie, M. & Siemann, E. (1997). The Influence of Functional Diversity and Composition on Ecosystem Processes. Science, 277, 1300–1302.

Urgoiti, J., Messier, C., Keeton, W.S., Reich, P.B., Gravel, D. & Paquette, A. (2022). No complementarity no gain—Net diversity effects on tree productivity occur once complementarity emerges during early stand development. Ecol. Lett., 25, 851–862.

Waterer, J.G. & Vessey, J.K. (1993). Effect of low static nitrate concentrations on mineral nitrogen uptake, nodulation, and nitrogen fixation in field pea. J. Plant Nutr., 16, 1775–1789.

Wen, Z., White, P.J., Shen, J. & Lambers, H. (2022). Linking root exudation to belowground economic traits for resource acquisition. New Phytol., 233, 1620–1635.

Xie, Z., Yu, Z., Li, Y., Wang, G., Liu, X., Tang, C., et al. (2022). Soil microbial metabolism on carbon and nitrogen transformation links the crop-residue contribution to soil organic carbon. Npj Biofilms Microbiomes, 8, 14.

Yang, C.-X., Chen, S.-J., Hong, X.-Y., Wang, L.-Z., Wu, H.-M., Tang, Y.-Y., et al. (2025). Plant exudates-driven microbiome recruitment and assembly facilitates plant health management. FEMS Microbiol. Rev., 49, fuaf008.

Zhou, Y., Zhu, H., Fu, S. & Yao, Q. (2017). Metagenomic evidence of stronger effect of stylo (legume) than bahiagrass (grass) on taxonomic and functional profiles of the soil microbial community. Sci. Rep., 7, 10195.

Zhu, S., Vivanco, J.M. & Manter, D.K. (2016). Nitrogen fertilizer rate affects root exudation, the rhizosphere microbiome and nitrogen-use-efficiency of maize. Appl. Soil Ecol., 107, 324–333.

