## Supplementary Information for "Active and total rhizosphere microbiomes of plant mixtures diverge from additive expectations, depending on nitrogen and plant composition"

#### Supplemental Methods

*Greenhouse experiment:* We grew crimson clover (legume, Trifolium incarnatum), canola (brassica, Brassica napus), and triticale (grass, Triticosecale) in monocultures, in pairwise combinations, and in three-species mixtures with and without nitrogen added in 4 randomized blocks (Figure 1, 7 plant treatments x 2 nitrogen levels x 6 replicates, N = 84). All pots contained 6 plants: 3 of each species in pairwise combinations and 2 of each species in pairwise mixtures. We refer to treatments by plant functional groups (legume, brassica, grass) rather than by species names. Nitrogen-fertilized treatments received 200 ppm aqueous ammonium nitrate every other week, and 300 ppm in the last week, while the unfertilized treatments received only tap water. We used field soil from a Russell E. Larson Agricultural Research Center at Rock Springs, PA (40.7215, −77.9275), which had a previous history of a cover crop mixture (field pea, crimson clover, triticale, canola, forage radish, and oats) grown every 3rd year on the plot for the past 12 years, in rotation with corn, soy, and a cool-season cereal (Finney, White and Kaye, 2016; Hosler et al., 2025). Soil samples collected before the start of the experiment were air-dried for 4 weeks to measure baseline nutrient levels (Table S1). To improve drainage, the field soil was amended with #1 Q-ROK sand (US Silica) and vermiculite in a 5:4:1 mixture by volume. To allow measurement of treatment effects on weed germination, 400 foxtail (Setaria faberi) and 500 pigweed (Amaranthus powellii) seeds were mixed into each pot prior to sowing of the focal seeds. Plants were grown for 12 weeks prior to measuring microbial activity, collecting rhizosphere samples, harvesting weed seeds, and measuring above- and below-ground biomass, to allow ample time for treatments to influence the microbial community. Four soil-only control pots were included to measure microbial community composition and activity without plant influence.

*Plant Biomass:* Aboveground and belowground biomass samples were separated by species, oven-dried for 1 week at 55°C, and weighed. Roots that grew into the cotton plugs at the bottom of the pots were dried and weighed separately because they could not unambiguously be divided by species. This fraction represented, on average, less than 10% of the total weight.

*Nitrogen fixation estimation using nitrogen stable isotopes:* To measure the proportion of plant nitrogen derived from biological nitrogen fixation (as opposed to chemical fertilizers), we used the natural abundance method. We collected triticale monoculture shoot tissue and legume shoot tissue from all cover crop treatments with legumes. The collected shoot tissue was oven-dried, weighed, and ground in a Udy vacuum cyclone with a 1mm screen. From the ground tissue, 3.25mg was tin-packed and submitted to Cornell University Stable Isotope Laboratory for $\delta^{15}N$ combustion analysis. We then used the following equation to calculate the percentage of nitrogen derived from the atmosphere (%Ndfa; Blesh et al. 2019, Pinto et al. 2021):

$$\%Ndfa = 100[(\delta^{15}N_{ref} - \delta^{15}N_{leg}) / (\delta^{15}N_{ref} - B)]$$

where $\delta^{15}N_{ref}$ is the $^{15}N$ relative natural abundance of the reference non-legume plant. We used triticale grown in monoculture and averaged across our six monoculture pots to estimate standard background $\delta^{15}N$. $\delta^{15}N_{leg}$ is each legume sample's $^{15}N$ relative natural abundance, and B is the $\delta^{15}N$ relative natural abundance of the legume when it is grown in a nitrogen-free medium. Blesh et al. (2019) calculated B as -1.55 for crimson clover. Each % nitrogen from biological nitrogen fixation (%Ndfa) was calculated for each sample, and technical replicates were averaged to estimate a value for each pot. With %Ndfa per pot calculated, we quantified BNF (Biological Nitrogen Fixation) using the following equation (Pinto et al., 2021):

BNF (g $pot^{-1}$) = legume biomass (g $pot^{-1}$) • (plant N concentration (%) / 100) • (%Ndfa / 100)

Here, we calculated BNF in grams per pot by multiplying the total legume stem biomass by the proportion of plant N concentration, and multiplying that by the previously calculated proportion of atmospheric-derived N. Because pots had different numbers of legume plants per treatment, we divided the treatment value by the number of legume plants per pot to obtain, for each pot BNF in grams per legume.

*Seed germination:* To determine seed germination in each treatment, we first added 400 foxtail (*Setaria faberi*) and 500 pigweed (*Amaranthus powellii*) seeds to each pot. At the end of the experiment, we collected weed seeds using sieves with openings ranging from 1 mm to 600 µm, then performed a germination test on a subset of 50 seeds per pot from nitrogen fertilized pots only. The weed seeds were placed on germination paper with 400 ppm gibberellic acid solution in parafilm-sealed petri dishes in an incubator at 30°C and allowed to germinate for 12 days with periodic rehydration. Dishes were checked every 3 days, germinated seeds were counted and removed, and the dishes' positions in the incubator were rearranged.

*Expected values for plant traits:* For above- and below-ground biomass, we compare measured values across treatments with expected values derived from monocultures. This approach allows us to compare values between monocultures and mixtures and is analogous to using the total relative yield (De Witt, 1960; Fowler, 1982). For each replicate, expected values in mixtures were calculated by multiplying the measured biomass in the monoculture by the planted ratio in the mixtures (Fowler, 1982). For example, for replicate 1, we predicted the shoot biomass for the legume grass mixtures by:

*Expected biomass LG = (L biomass + G biomass)* • (1/2)

This approach yielded expected values for each rep of shoot and root biomass, which could be compared with the measured values. This was extended to three-species mixtures, in which the expected value for a three-species mixture was one-third the value from each species monoculture.

*Statistical analysis of biomass, nitrogen, and seed germination:* All statistical analysis was conducted in R (version 4.5.1). First, in a linear model, we tested for differences in shoot and root biomass, percent nitrogen from fixation, and nitrogen fixed per legume using the "aov" function (stats package, R Core Team, 2025). For all models, block effect was included as a fixed effect, and nitrogen level was included as an interaction effect. We confirmed normality with the Kolmogorov-Smirnov test and visual assessment of qq plots of the residuals. We conducted Tukey post hoc tests using estimated marginal means from the emmeans package (Russell V. Lenth and Julia Piaskowski, 2026). We compared expected vs. measured shoot and root biomass in mixtures using separate ANOVAs for each mixture with rep as a fixed effect and nitrogen level as an interaction effect. We used a similar approach to analyze weed seed germination (foxtail and pigweed) with a binomial generalized linear model and a Tukey-Sidak post hoc test (model formula, nonviable: viable weed seeds ~ treatment + block). Then we compared the expected vs measured using a binomial model (nonviable: viable weed seeds ~ treatment + block). Because weed seed germination tests were only done on seeds from nitrogen fertilized pots, nitrogen was not included in the model.

*BONCAT click chemistry and flow cytometry:* Then, rhizosphere/glycerol aliquots for activity assessment and sorting were thawed and prepped for FACS (Fluorescence-Activated Cell Sorting). We removed excess soil by centrifuging the rhizosphere slurry at 500 x g for 5 minutes, retaining the cell-containing supernatant. To pellet the cells, we centrifuged the cell-containing mixture at 14,000 x g for 5 minutes and removed the supernatant. Then we resuspended the cells in 300 µL of PBS. Next, HPG-containing cells were fluorescently labeled with the FAM picolyl azide dye (Click Chemistry Tools; Ex/Em 490/510) via a click chemistry reaction adapted from Reichardt (2020). Click reaction mix consisting of 5 µl copper sulfate (200 µM final concentration), 10 µl tris-hydroxypropyltriazolylmethylamine (1000 µM final concentration), 3.3 µl FAM picolyl azide dye (10 µM final concentration), buffered with 50 µl of 5 mM sodium ascorbate in 1M PBS, 50 µl of 5 mM aminoguanidine HCl in 1M PBS, and 380 µl of 1M PBS. We added 200 µL of the click master mix directly to the cell mixture and incubated the cells for 1 hour at 27°C while shaking at 200 rpm. The click reaction was stopped by pelleting cells and washing the pellet 3 times with PBS. Finally, we added SYTO59 DNA counterstain (Invitrogen, Ex/Em 622/645 nm) at a final concentration of 0.5 µM.

We initially determined that microbial activity was present in all treatments using the BD Fortessa Cytometer at the Huck Flow Cytometry Facility. Because microbial activity was close to the detection limit across multiple replicates of the unfertilized treatments, we isolated active microbial cells only from the nitrogen-fertilized treatments to ensure sufficient sample sizes for comparison. We sorted active microbial cells from all seven cover crop mixtures and soil-only controls and measured microbial activity on the Beckman Coulter MoFlo Astrios EQ Cell Sorter. On both instruments, the threshold size was set to above 0.2 µm determined by Flow Cytometry Sub-Micron Particle Size Reference Kit Beads (Invitrogen), and dictation was set up to capture the FAM picolyl azide dye (Ex/Em 490/510 nm) in the green channel of a 488 nm blue laser and the SYTO59 DNA counterstain (Ex/Em 622/645 nm) in the red channel of a 630 nm red laser. Viable cells were identified by drawing a SYTO59-positive gate against an unstained control. Active cells were gated from the SYTO59-positive events using a nested gating strategy. Active cells were determined by BONCAT FAM picolyl azide-positive events with a water-incubated clicked control to account for background fluorescence from the click reaction. We measured the percentage of active cells for 3 minutes per sample, with a false-positive rate capped at 1%. Then, we collected all active microbial cells from a known volume of the cell mixture and used that value to estimate the number of active microbial cells per gram of rhizosphere soil. In addition to the active microbial cells, we collected inactive microbial cells in all nitrogen-fertilized treatments. On average, we collected 65k active cells per sample, with a minimum of 26k.

*Statistical analysis of microbiome activity, composition, and diversity:* Differences in the percentage of active microbial cells between treatments were analyzed using a binomial model (percent active ~ treatment + block). Pairwise differences were determined with a Tukey corrected post hoc test. Differences in the number of active cells per gram of rhizosphere soil were analyzed using a negative binomial model, appropriate for count data, and pairwise differences were assessed using a Tukey corrected post hoc test (counts ~ treatment + block).

We calculated microbial alpha diversity, including Shannon diversity and species richness, using Phyloseq (McMurdie & Holmes, 2013). Pilou's Evenness was calculated using the formula $J = H/\ln(S)$, where J is the Pilou's Evenness, H is Shannon diversity, and S is the number of species. For all alpha diversity metrics from the total rhizosphere microbial community, the full model was diversity ~ N*treatment*block. However, we didn't detect significant effects of Block, so we dropped it from the model (final model: Diversity ~ treatment * N). Differences between treatments were determined with ANOVA and a post hoc Tukey test. We carried out the same analysis for the active community but did not test for a nitrogen effect, as we measured the active community only in nitrogen-fertilized treatments (final model: diversity ~ treatment +block ).

In our analysis of microbial community composition, we first filtered out rare taxa with fewer than 5 reads per sample on average (i.e., less than 0.0001% abundance). We then calculated Bray-Curtis dissimilarity matrices and ordinated them using Canonical Analysis of Principal Coordinates (CAP) from the R package vegan (Oksanen, Jari et al., 2022). For the total rhizosphere samples, we ran a CAP model to test the effects of nitrogen and treatment, with block as a fixed effect (distance matrix ~ treatment*nitrogen*block). Within the total rhizosphere community, we found a marginal interaction between nitrogen and treatment ($p = 0.05$). We then subset the total DNA into nitrogen-fertilized and unfertilized and examined how treatment effects differed between the nitrogen treatments. We made pairwise comparisons between treatments using the R package BiodiversityR (Roeland Kindt & Richard Coe, 2005). For the active and inactive samples, we ran a full CAP model to assess the effects of treatment and fraction (active vs inactive; formula: distance matrix ~ treatment*fraction + block), followed by a pairwise comparison of the treatments for both the inactive and active fraction using a reduced model with block as a random effect (distance matrix ~ treatment+ condition(block)). In both the total and active communities, we assessed the contribution of each plant functional group to microbial composition in a CAP, in which the presence or absence of plant functional groups is the explanatory variable (distance matrix ~ Legume * Grass * Brassica + block).

We identified key taxa driving differences between treatments by evaluating which ASVs had the longest vectors on CAP1 and CAP2 in both the nitrogen-fertilized active and total communities.

We calculated their distances from the origin and retained the 20 ASVs with the largest distances. To determine statistical differences in the abundance of these ASVs between treatments, we used DESeq2 (Love, Huber, and Anders, 2014). We ran a DEseq2 analysis with the presence vs. absence of each plant functional group and their interactions as predictors, to explore the contributions of each plant species to the relative abundance of key taxa (Abundance ~ L*G*B). Soil-only pots were included in the dataset for DEseq2 analysis to provide a baseline when all plants are absent.

*Calculating and testing for expected microbial activity and microbiome composition:* We calculated the expected microbial activity (active cells/g rhizosphere and % active cells) in a mixture from the monocultures. For each replicate, expected values were calculated by multiplying each monoculture's value by the measured percentage root biomass for that species, then summing the results across monocultures. For example, for active cells/g in replicate one of the legume-grass mixture, the formula used is shown below:

Active cells/g LG = (active cells/g L) • (% root biomass of L in LG) +

(active cells/g G) • (% root biomass of G in LG)

We compared the active cells/g between expected and measured values using a separate ANOVA for each mixture. We compared the percentage of active cells between mixtures and monocultures using a binomial model for each mixture.

We calculated the expected ASV abundances in mixtures using monoculture data. We used the same rare-taxon filtering approach as for community composition (excluding taxa with fewer than 5 reads per sample). For each replicate, expected values were calculated by multiplying the ASV abundance from each monoculture by the measured percentage root biomass of that species and summing the resulting abundance across monocultures. For example, for each ASV in one replicate of the legume grass mixture, the formula used is shown below.

ASV abundance LG = (ASV abundance L) • (% root biomass of L in LG) +

(ASV abundance in G) • ( % root biomass of G in LG)

We compared the abundance and presence of microbial ASVs between expected values from monocultures and measured values from mixtures. For both the total and active microbiomes, we applied a Centered Log-Ratio (CLR) transformation. A CLR transformation converts compositional data into a standard Euclidean space or Aitchison distance, which we then ordinated with a PCA. Using a PERMANOVA, we tested whether there was a significant difference between the expected and measured communities for each cover crop mixture. We then compared the measured microbial composition of pairwise mixtures with that of their respective monocultures using a scale vector projection. We projected the composition data for each pairwise mixture onto the vector connecting the monoculture centroids. We scaled values to -1 to 1, with 0 representing the expected value. We extended this method to the three-species mixture by calculating the position of the mixture along the vector connecting the centroids of each monoculture. We did not normalize the three-species mixture to -1 to 1; instead, we visualized the raw values from 0 to 1 with ggtern (Nicholas E. Hamilton & Michael Ferry, 2018), since negative numbers cannot be plotted in ternary diagrams, which rely on barycentric coordinates.

### Supplemental Figures

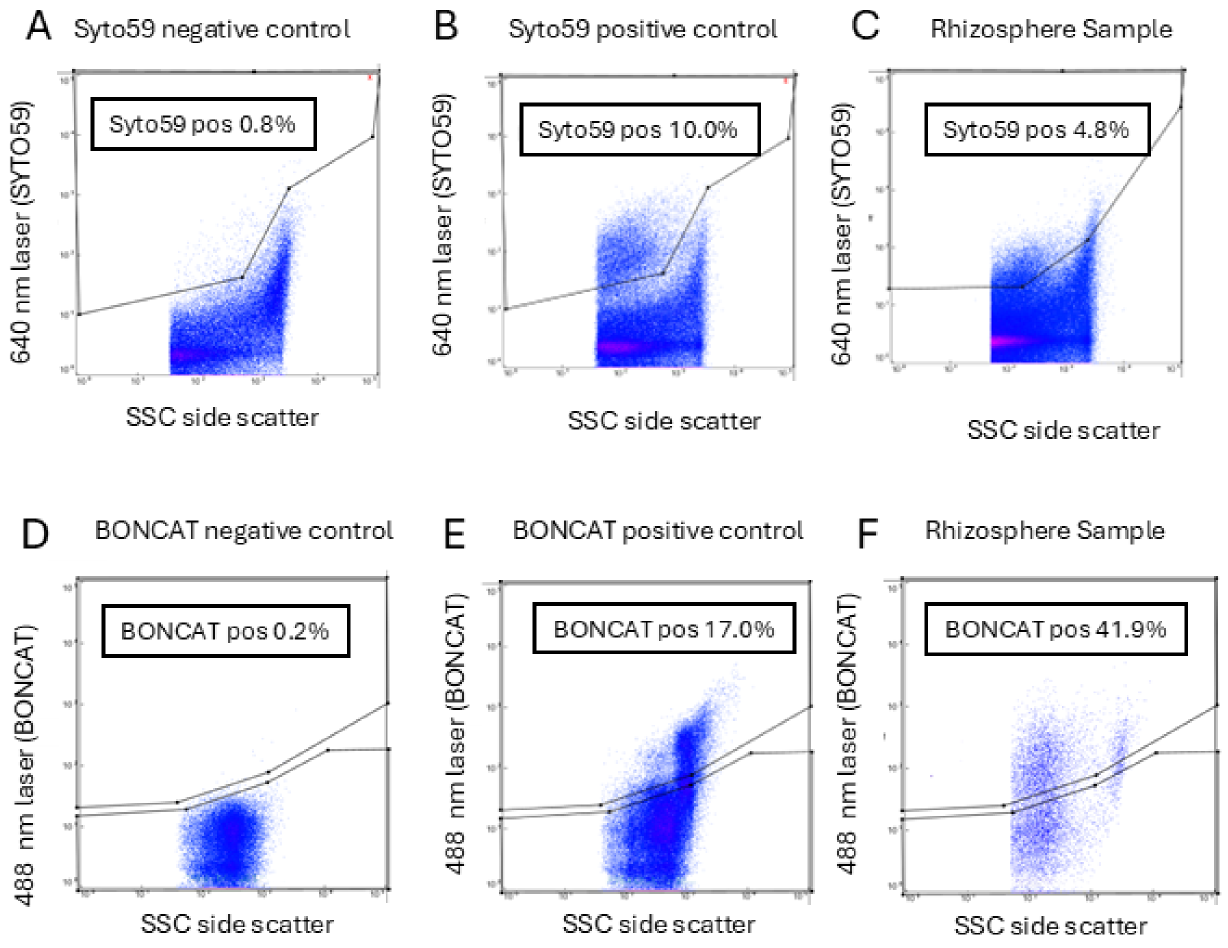


*Supplemental Figure S1:* Examples of gating of rhizosphere samples on a flow cytometer to define bacterial cells and active cells. (A–C). Nucleic acid staining for cell identification from soil particles. Flow cytometry data showing side scatter (SSC) versus SYTO 59 fluorescence intensity (640-nm excitation), both on a log scale. Representative gates delineate SYTO 59-positive events, which are identified as bacterial cells over soil debris by fluorescence and size. We report the percentage of events that are bacterial cells (Syto59 pos) for (A) the unstained negative control (0.8%), (B) the SYTO 59-stained positive control (10.0%), and (C) the stained rhizosphere sample (10.0%). (D–F) Bioorthogonal non-canonical amino acid tagging (BONCAT) for active cell detection: in a nested gating strategy, only bacterial cells are shown (SYTO 59 positive). Flow cytometry data is plotted, displaying SSC versus BONCAT dye fluorescence intensity (488 nm excitation), both on a log scale. Flow cytometry gates define translationally active cells within the total bacterial cells for the (D) negative control (unlabeled/click-reaction negative control, 0.2%), (E) positive control (17.0%), and (F) experimental rhizosphere sample

(41.9% BONCAT-positive). Additional flow cytometry scatter plots on Figshare DOI: 10.6084/m9.figshare.33292464

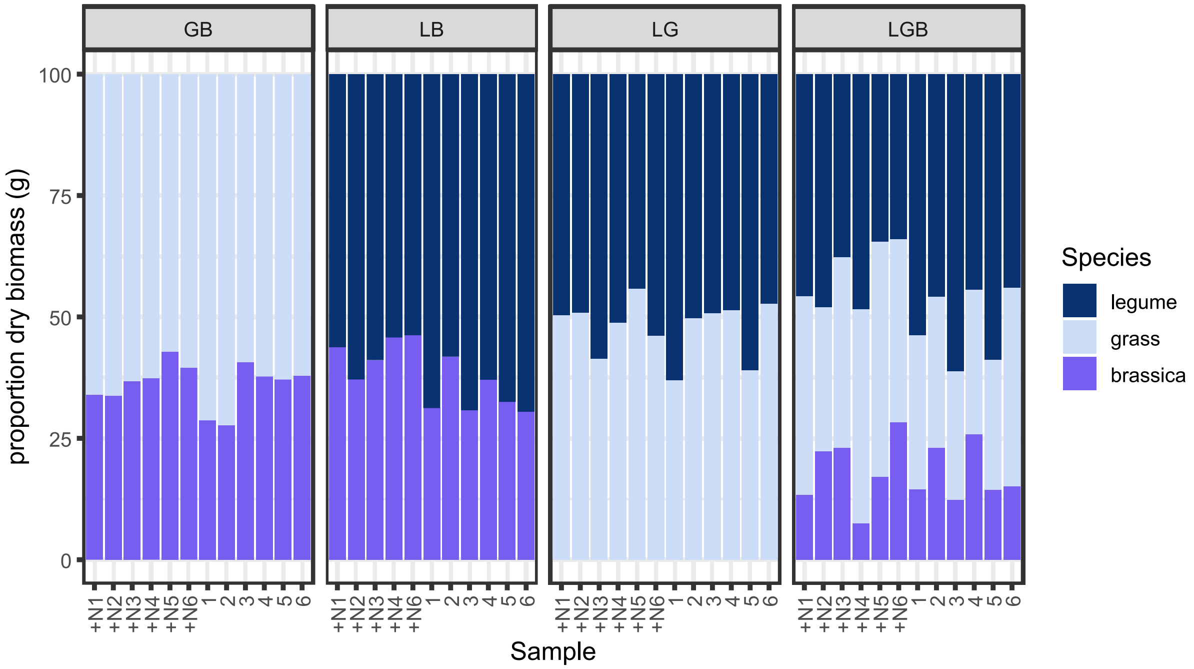


*Supplemental Figure S2:* Proportion of root dry biomass belonging to each species in each of the experimental mixture pots at harvest time. Monocultures are excluded because they consist of a single species. Data are grouped by treatment, where GB is grass-brassica mixture, LB is legume-brassica mixture, LG is legume-grass mixture, and LGB is legume-grass-brassica mixture. In the x-axis labels, +N indicates nitrogen-fertilized pots, and the number indicates the replicate.

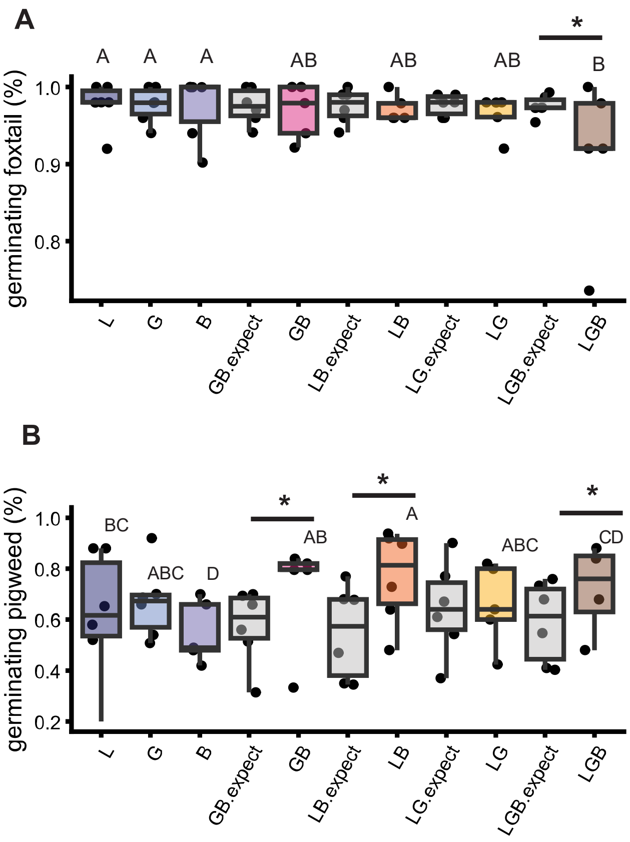


*Supplemental Figure S3*: Seed germination of A) foxtail and B) pigweed seed. Letters indicate pairwise differences between Treatments in a binomial model that included experimental data. In addition, we calculated expected values in each mixture (shown in gray and appended with "expect"). Asterisks indicate significant differences between the measured values for a given mixture treatment and the additive expectations.

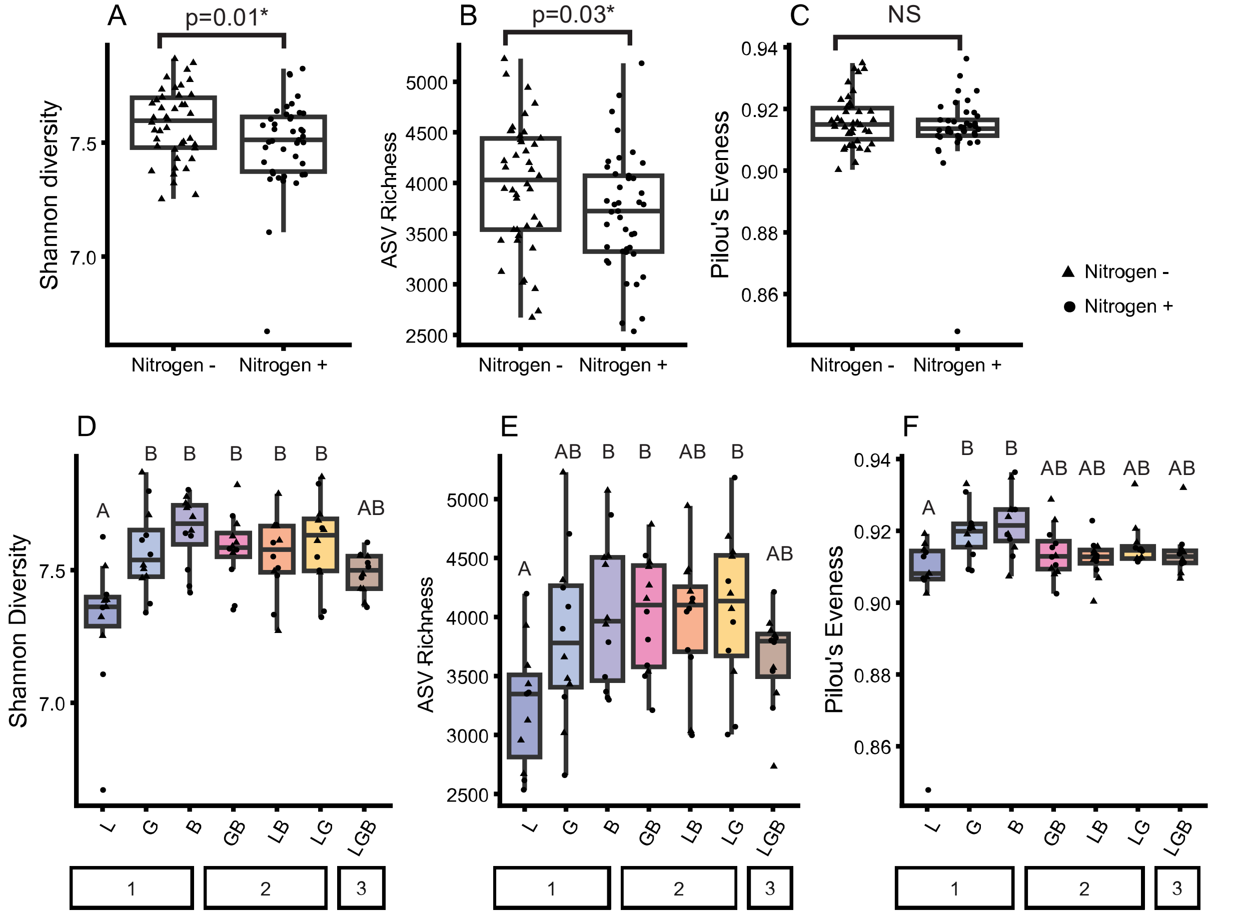


*Supplemental Figure S4*: Effect of nitrogen addition (A-C) and plant treatments (D-E) on alpha diversity of the total rhizosphere microbiome measured by 16S rRNA gene amplicon sequencing. A) Shannon diversity index, B) ASV richness, and C) Pilou's evenness. (A-C) P-values show significant differences between nitrogen levels in a linear model. D) Shannon diversity index, E) ASV richness, and F) Pilou's evenness. (D-F) Letters indicate significant differences in a Tukey post hoc test (linear model: Diversity metric ~ Treatment + Block). Treatments are grouped by the number of species. All measurements are calculated on rarefied amplicon sequence variants (ASVs).

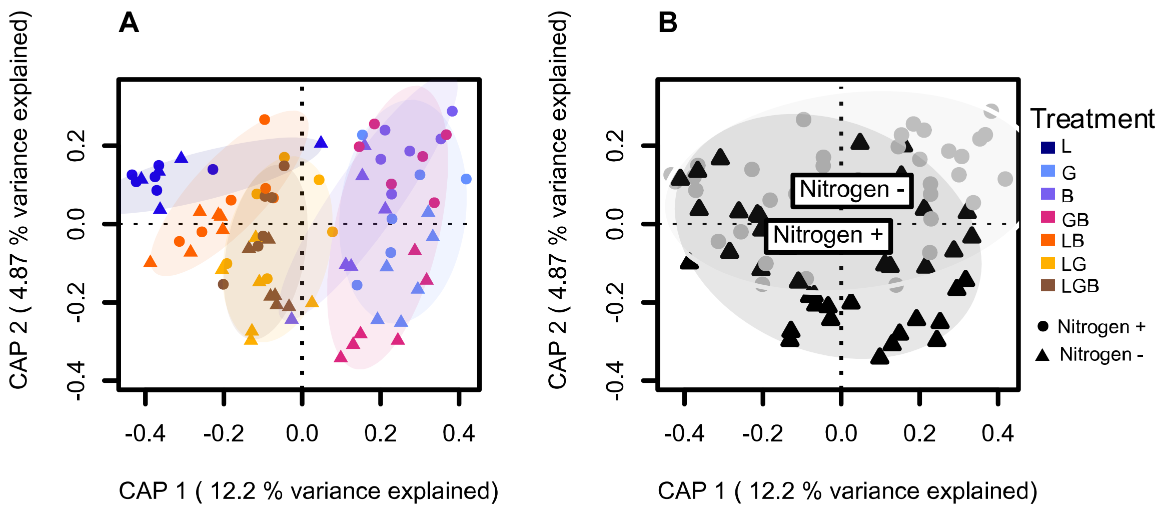


*Supplemental Figure S5:* Nitrogen amendment significantly shifts total rhizosphere microbial community structure. A) CAP visualization of the effect of both treatment and nitrogen on Bray-Curtis dissimilarities for rhizosphere total microbiome, colored by plant community, model results in table S26. B) depicts the same CAP analysis as A, but with the data coded to highlight Nitrogen amendment treatment (black triangles are N- and grey circles are N+).

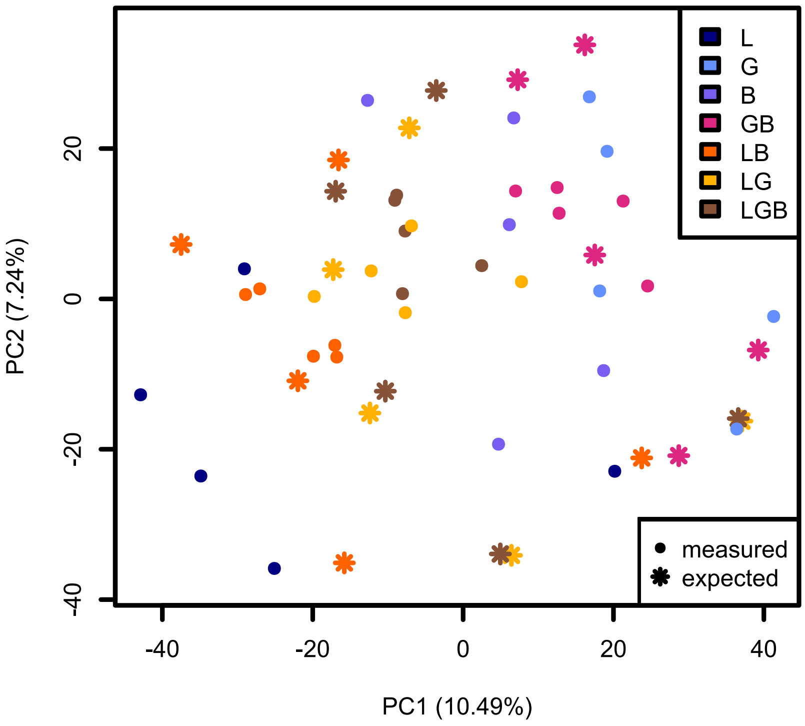


*Supplemental Figure S6:* PCA of Aitchison distances for the expected (stars) vs measured (dots) total microbial community, see supplemental methods for details on expected microbiome calculations. Because replicate 2 could not be amplified in the Legume monoculture, it was excluded from calculations for all mixtures, resulting in only 5 replicates shown. Aitchison distances were used to project the composition of mixtures onto a vector between the centroids of the monocultures, as shown in Figure 3.

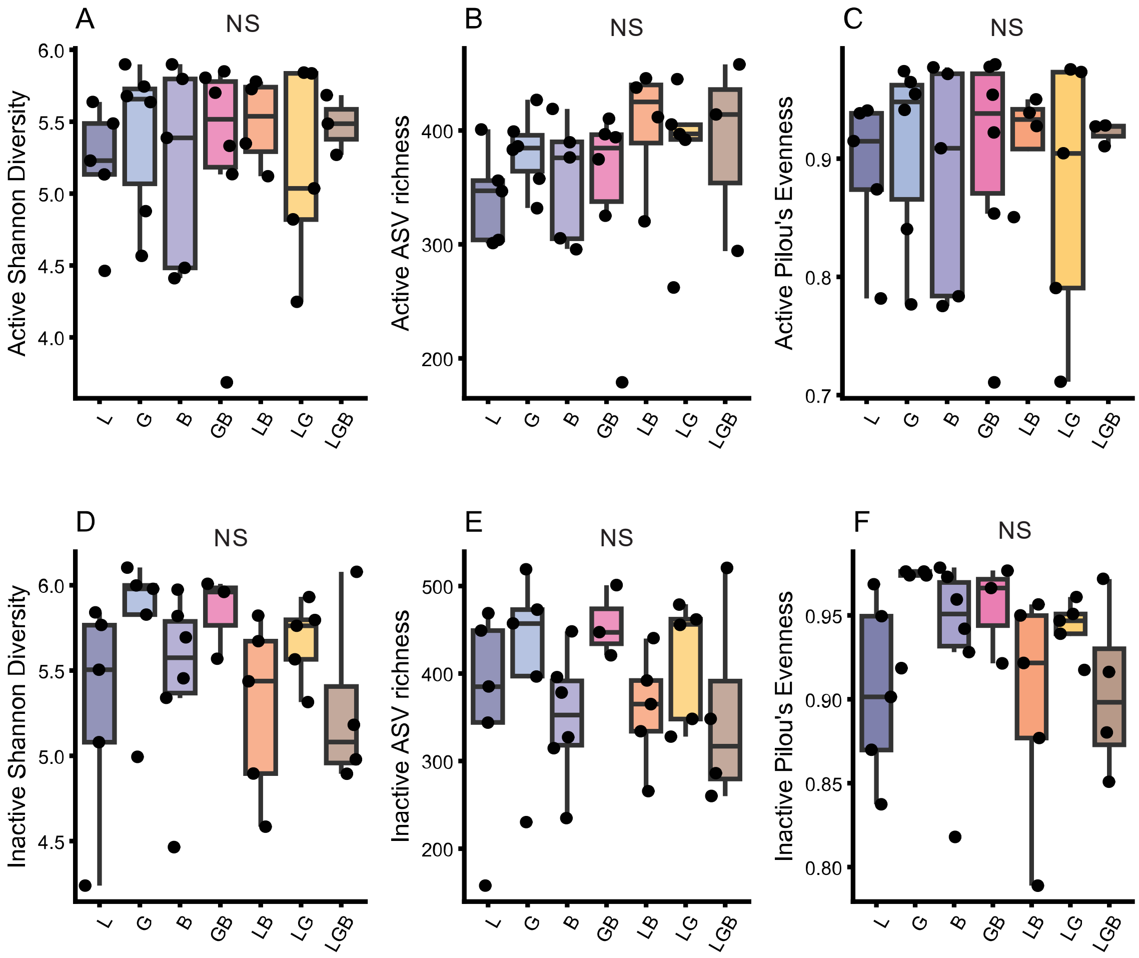


*Supplemental Figure S7:* Microbiome alpha diversity in the active (top) and inactive (bottom) cell fraction in the rhizosphere, including A,D) Shannon diversity B,E) species richness and C,F) Pilou's evenness. No significant treatment differences were detected using linear models followed by ANOVA and Tukey post hoc tests (Diversity metric ~ Treatment + block).

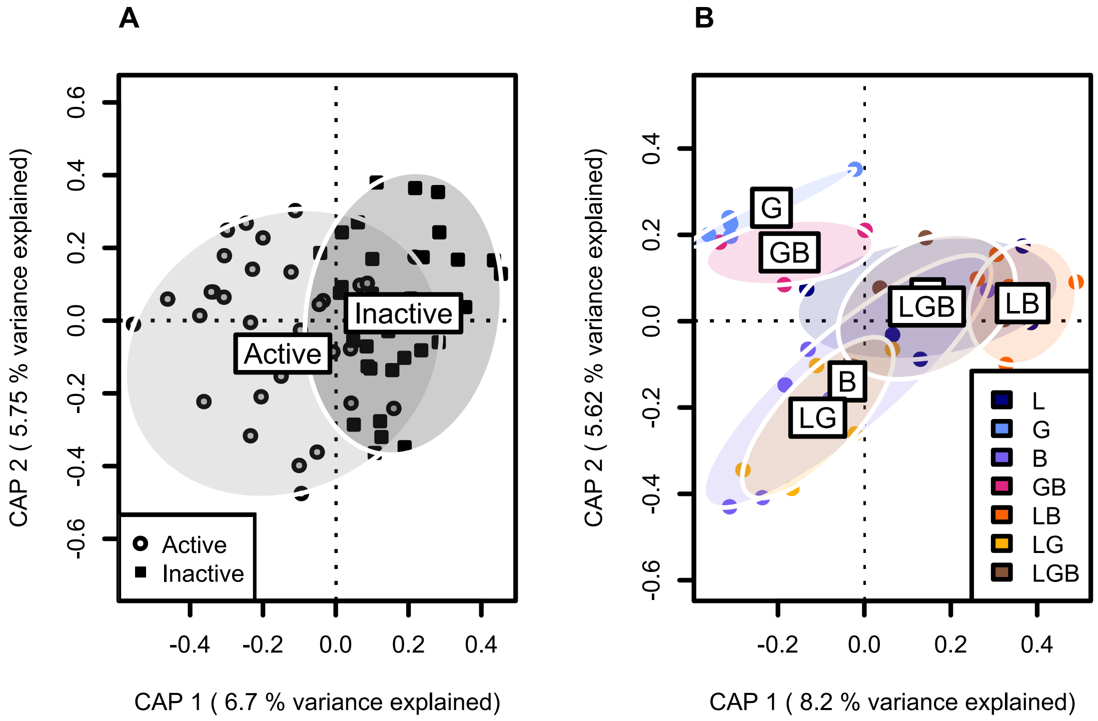


*Supplemental Figure S7:* Active and inactive microbial communities are distinct and impacted by plant treatment. (A) CAP visualization of the effect of treatment, fraction (active vs. inactive), and block on Bray-Curtis dissimilarities for the sorted rhizosphere microbial community. Model results are in Table S39. B) CAP visualization of  Bray-Curtis dissimilarities for the inactive rhizosphere microbiome of the effects of treatment and block. CAP ordination of Ellipses represent 95% confidence intervals. Pairwise differences calculated in a permutation test are shown in Table S39.

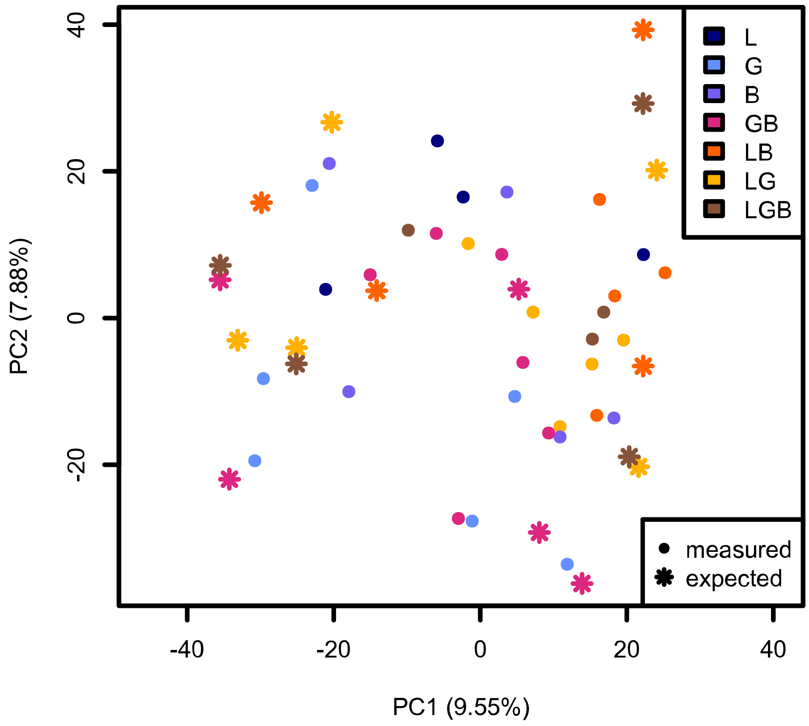


*Supplemental Figure S8:* PCA of Aitchison distances for the expected (stars) vs measured (dots) active microbial community. Aitchison distances are centered, log-transformed relative abundances from the measured and predicted communities; see supplemental methods for details on expected microbiome calculations. Aitchison distances were used to project the composition of mixtures onto a vector between the centroids of the monocultures, as shown in Figure 4.

### Supplemental Tables

*Table S1:* Nutrient report on field soil and sand/vermiculite amended field soils. Soils measured by Penn State Ag analytical labs.

| *Nutrient tested* | *Field soil* | *Sand/vermiculite amended field soil* |
|---|---|---|
| *Soil pH* | 6.9 | 6.8 |
| *Phosphorus (P)* | 19 ppm | 15 ppm |
| *Potassium (K)* | 152 ppm | 90 ppm |
| *Magnesium (Mg)* | 107 ppm | 55 ppm |

*Table S2 :* ANOVA results for the model on shoot biomass (grams dry weight). Model: shoot biomass ~ treatment* nitrogen +block. Data shown in Figure 2.

| *Term* | *df* | *Sum Sq* | *Mean Sq* | *F value* | *p value* |
|---|---|---|---|---|---|
| *Treatment* | 6 | 503.3 | 83.89 | 79.336 | < 2e-16* |
| *Nitrogen* | 1 | 4.4 | 4.41 | 4.169 | 0.04506* |
| *Treatment* Nitrogen* | 6 | 19.4 | 3.24 | 3.061 | 0.01036* |
| *Block* | 1 | 8.3 | 8.26 | 7.816 | 0.00672* |
| *Residuals* | 68 | 71.9 | 1.06 | | |

*Significant p<0.05

*Table S3:* Pairwise comparisons of shoot biomass across treatments within nitrogen levels. Estimated marginal means (EMMs) and standard errors were calculated from an ANOVA model including treatment, nitrogen level, and block effects (n= 83). Pairwise differences between treatments were evaluated separately within each nitrogen level using Tukey's HSD test. The results below are from samples without added nitrogen. Data shown in Figure 2.

| *contrast* | *estimate* | *SE* | *df* | *t ratio* | *p value* |
|---|---|---|---|---|---|
| *L - G* | 8.5983 | 0.594 | 68 | 14.483 | <0.0001* |
| *L - B* | 7.2717 | 0.594 | 68 | 12.249 | <0.0001* |
| *L - GB* | 8.2482 | 0.595 | 68 | 13.872 | <0.0001* |
| *L - LB* | 3.4884 | 0.594 | 68 | 5.874 | <0.0001* |
| *L - LG* | 3.4717 | 0.594 | 68 | 5.848 | <0.0001* |
| *L - LGB* | 4.2634 | 0.594 | 68 | 7.179 | <0.0001* |
| *G - B* | -1.3267 | 0.594 | 68 | -2.235 | 0.2913 |
| *G - GB* | -0.3501 | 0.595 | 68 | -0.589 | 0.9969 |
| *G - LB* | -5.11 | 0.594 | 68 | -8.604 | <0.0001* |
| *G - LG* | -5.1267 | 0.594 | 68 | -8.635 | <0.0001* |
| *G - LGB* | -4.335 | 0.594 | 68 | -7.299 | <0.0001* |
| *B - GB* | 0.9766 | 0.595 | 68 | 1.642 | 0.6558 |
| *B - LB* | -3.7833 | 0.594 | 68 | -6.37 | <0.0001* |
| *B - LG* | -3.8 | 0.594 | 68 | -6.401 | <0.0001* |
| *B - LGB* | -3.0083 | 0.594 | 68 | -5.065 | <0.0001* |
| *GB - LB* | -4.7599 | 0.596 | 68 | -7.989 | <0.0001* |
| *GB - LG* | -4.7766 | 0.595 | 68 | -8.033 | <0.0001* |
| *GB - LGB* | -3.9849 | 0.596 | 68 | -6.689 | <0.0001* |
| *LB - LG* | -0.0167 | 0.594 | 68 | -0.028 | 1 |
| *LB - LGB* | 0.775 | 0.594 | 68 | 1.305 | 0.8471 |
| *LG - LGB* | 0.7917 | 0.594 | 68 | 1.333 | 0.834 |

*Significant p<0.05

*Table S4:* Pairwise comparisons of shoot biomass across treatments within nitrogen levels. Estimated marginal means (EMMs) and standard errors were calculated from an ANOVA model including fixed effects of treatment, nitrogen level, and block (n= 83). Pairwise differences between treatments were evaluated separately within each nitrogen level using Tukey's HSD test. The results below are from samples with added nitrogen. Data shown in Figure 2

| *contrast* | *estimate* | *SE* | *df* | *t ratio* | *p value* |
|---|---|---|---|---|---|
| *L - G* | 6.2984 | 0.595 | 68 | 10.593 | <0.0001* |
| *L - B* | 4.5768 | 0.595 | 68 | 7.697 | <0.0001* |
| *L - GB* | 5.3584 | 0.594 | 68 | 9.022 | <0.0001* |
| *L - LB* | 1.7943 | 0.623 | 68 | 2.882 | 0.0742 |
| *L - LG* | 2.45 | 0.594 | 68 | 4.125 | 0.0019* |
| *L - LGB* | 3.1802 | 0.597 | 68 | 5.323 | <0.0001* |
| *G - B* | -1.7217 | 0.594 | 68 | -2.9 | 0.0709 |
| *G - GB* | -0.94 | 0.594 | 68 | -1.583 | 0.6935 |
| *G - LB* | -4.5041 | 0.623 | 68 | -7.225 | <0.0001* |
| *G - LG* | -3.8484 | 0.594 | 68 | -6.48 | <0.0001* |
| *G - LGB* | -3.1182 | 0.595 | 68 | -5.244 | <0.0001* |
| *B - GB* | 0.7816 | 0.594 | 68 | 1.316 | 0.8421 |
| *B - LB* | -2.7824 | 0.623 | 68 | -4.463 | 0.0006* |
| *B - LG* | -2.1267 | 0.594 | 68 | -3.581 | 0.0109* |
| *B - LGB* | -1.3966 | 0.595 | 68 | -2.349 | 0.2367 |
| *GB - LB* | -3.564 | 0.623 | 68 | -5.723 | <0.0001* |
| *GB - LG* | -2.9083 | 0.594 | 68 | -4.899 | 0.0001* |
| *GB - LGB* | -2.1782 | 0.596 | 68 | -3.656 | 0.0086* |
| *LB - LG* | 0.6557 | 0.623 | 68 | 1.053 | 0.9394 |
| *LB - LGB* | 1.3858 | 0.626 | 68 | 2.214 | 0.3017 |
| *LG - LGB* | 0.7301 | 0.596 | 68 | 1.226 | 0.8818 |

*Significant p<0.05

*Table S5:* Pairwise differences between expected and measured values in a one-way ANOVA for shoot biomass (g dry weight). Model: shoot biomass ~ measurement type*nitrogen. There was no significant interaction between nitrogen and measurement type. Data shown in Figure 2

| *Treatment* | *contrast* | *df (residual)* | *Sum sq (treatment)* | *Sum sq (residual)* | *F value* | *p value* |
|---|---|---|---|---|---|---|
| *GB* | expected vs measured | 20 | 0.0104 | 1.89997 | 0.1097 | 0.7440 |
| *LB* | expected vs measured | 19 | 0.4706 | 19.3699 | 0.4616 | 0.5051 |
| *LG* | expected vs measured | 20 | 3.4961 | 25.4437 | 2.7481 | 0.1130 |
| *LGB* | expected vs measured | 20 | 2.4897 | 12.8023 | 3.8895 | 0.0625 |

*Table S6:* ANOVA results for the model on root biomass (grams dry weight). Model: root biomass ~ treatment*nitrogen +block. Data shown in Figure 2.

| *Term* | *df* | *Sum Sq* | *Mean Sq* | *F value* | *p value* |
|---|---|---|---|---|---|
| *Treatment* | 6 | 33.11 | 5.518 | 28.491 | < 2e-16* |
| *Nitrogen* | 1 | 1.92 | 1.917 | 9.896 | 0.00246* |
| *Block* | 1 | 3.07 | 3.067 | 15.834 | 0.00017* |
| *Treatment* Nitrogen* | 6 | 2.7 | 0.451 | 2.327 | 0.04201* |
| *Residuals* | 68 | 13.17 | 0.194 | | |

*Significant p<0.05

*Table S7:* Pairwise comparisons of root biomass (grams dry weight) across treatments within nitrogen levels. Estimated marginal means (EMMs) and standard errors were calculated from an ANOVA model including fixed effects of treatment, nitrogen level, and block (n= 83). Pairwise differences between treatments were evaluated separately within each nitrogen level using Tukey's HSD test. The results below are from samples without added nitrogen (Nitrogen -). Data shown in Figure 2.

| *contrast* | *estimate* | *SE* | *df* | *t ratio* | *p value* |
|---|---|---|---|---|---|
| *L - G* | 1.09 | 0.254 | 68 | 4.29 | 0.0011* |
| *L - B* | 1.7683 | 0.254 | 68 | 6.96 | <0.0001* |
| *L - GB* | 1.4446 | 0.254 | 68 | 5.677 | <0.0001* |
| *L - LB* | 0.6885 | 0.254 | 68 | 2.709 | 0.1117 |
| *L - LG* | -0.2367 | 0.254 | 68 | -0.931 | 0.9661 |
| *L - LGB* | 0.1552 | 0.254 | 68 | 0.611 | 0.9963 |
| *G - B* | 0.6783 | 0.254 | 68 | 2.67 | 0.122 |
| *G - GB* | 0.3546 | 0.254 | 68 | 1.393 | 0.8035 |
| *G - LB* | -0.4015 | 0.254 | 68 | -1.579 | 0.6956 |
| *G - LG* | -1.3267 | 0.254 | 68 | -5.222 | <0.0001* |
| *G - LGB* | -0.9348 | 0.254 | 68 | -3.678 | 0.0081* |
| *B - GB* | -0.3238 | 0.254 | 68 | -1.272 | 0.8621 |
| *B - LB* | -1.0798 | 0.254 | 68 | -4.248 | 0.0013* |
| *B - LG* | -2.005 | 0.254 | 68 | -7.891 | <0.0001* |
| *B - LGB* | -1.6131 | 0.254 | 68 | -6.346 | <0.0001* |
| *GB - LB* | -0.756 | 0.255 | 68 | -2.965 | 0.0603 |
| *GB - LG* | -1.6812 | 0.254 | 68 | -6.607 | <0.0001* |
| *GB - LGB* | -1.2894 | 0.255 | 68 | -5.057 | <0.0001* |
| *LB - LG* | -0.9252 | 0.254 | 68 | -3.64 | 0.0091* |
| *LB - LGB* | -0.5333 | 0.254 | 68 | -2.099 | 0.365 |
| *LG - LGB* | 0.3919 | 0.254 | 68 | 1.542 | 0.7188 |

*Significant p<0.05

*Table S8:* Pairwise comparisons of root biomass (grams dry weight) across treatments within nitrogen levels. Estimated marginal means (EMMs) and standard errors were calculated from an ANOVA model including treatment, nitrogen level, and block effects (n= 83). Pairwise differences between treatments were evaluated separately within each nitrogen level using Tukey's HSD test. The results below are from samples with added nitrogen (Nitrogen +). Data shown in Figure 2.

| *contrast* | *estimate* | *SE* | *df* | *t ratio* | *p value* |
|---|---|---|---|---|---|
| *L - G* | -0.0129 | 0.254 | 68 | -0.051 | 1 |
| *L - B* | 1.1671 | 0.254 | 68 | 4.586 | 0.0004* |
| *L - GB* | 0.4952 | 0.254 | 68 | 1.948 | 0.4565 |
| *L - LB* | 0.3747 | 0.266 | 68 | 1.406 | 0.7967 |
| *L - LG* | -0.8831 | 0.254 | 68 | -3.474 | 0.0149* |
| *L - LGB* | -0.1225 | 0.256 | 68 | -0.479 | 0.999 |
| *G - B* | 1.18 | 0.254 | 68 | 4.644 | 0.0003* |
| *G - GB* | 0.5081 | 0.254 | 68 | 1.999 | 0.4248 |
| *G - LB* | 0.3876 | 0.267 | 68 | 1.453 | 0.771 |
| *G - LG* | -0.8702 | 0.254 | 68 | -3.424 | 0.0173* |
| *G - LGB* | -0.1096 | 0.254 | 68 | -0.431 | 0.9995 |
| *B - GB* | -0.6719 | 0.254 | 68 | -2.643 | 0.1294 |
| *B - LB* | -0.7924 | 0.267 | 68 | -2.97 | 0.0595* |
| *B - LG* | -2.0502 | 0.254 | 68 | -8.066 | <0.0001* |
| *B - LGB* | -1.2896 | 0.254 | 68 | -5.068 | <0.0001* |
| *GB - LB* | -0.1205 | 0.267 | 68 | -0.452 | 0.9993 |
| *GB - LG* | -1.3783 | 0.254 | 68 | -5.425 | <0.0001* |
| *GB - LGB* | -0.6177 | 0.255 | 68 | -2.423 | 0.2053 |
| *LB - LG* | -1.2578 | 0.267 | 68 | -4.719 | 0.0002* |
| *LB - LGB* | -0.4972 | 0.268 | 68 | -1.856 | 0.5159 |
| *LG - LGB* | 0.7606 | 0.255 | 68 | 2.983 | 0.0576* |

*Significant $p<0.05$

*Table S9:* Pairwise differences between expected and measured values in a one-way ANOVA for root biomass (g dry weight). Model: root biomass ~ measurement type*nitrogen. There was no significant interaction between nitrogen and measurement type. Data shown in Figure 2.

| *Treatment* | *contrast* | *df (residual)* | *Sum sq (treatment)* | *Sum sq (residual)* | *F value* | *p value* |
|---|---|---|---|---|---|---|
| *GB* | expected vs measured | 20 | 0.8513 | 4.7275 | 3.6013 | 0.07226 |
| *LB* | expected vs measured | 19 | 3.4768 | 7.8787 | 8.3845 | 0.00927* |
| *LG* | expected vs measured | 20 | 22.952 | 24.5703 | 18.6825 | 0.0003* |
| *LGB* | expected vs measured | 20 | 9.7538 | 4.5378 | 42.9892 | 2.177e-06* |

*Significant p<0.05

*Table S10:* ANOVA results on the main model for the percentage of nitrogen from biological nitrogen fixation (BNF). Model: percent nitrogen from BNF ~ treatment* nitrogen/block. Data shown in Figure 2.

| *Term* | *df* | *Sum Sq* | *Mean Sq* | *F value* | *p value* |
|---|---|---|---|---|---|
| *Treatment* | 3 | 1523.1 | 507.7 | 37.98 | 2.84E-12* |
| *Nitrogen* | 1 | 703 | 703 | 52.588 | 4.90E-09* |
| *Block* | 1 | 25.4 | 25.4 | 1.9 | 0.17501 |
| *Treatment* Nitrogen* | 3 | 349.2 | 116.4 | 8.709 | 0.00012* |
| *Residuals* | 44 | 588.2 | 13.4 | | |

*Significant p<0.05

*Table S11:* Pairwise comparisons of the percentage of nitrogen from biological nitrogen fixation (BNF ) across treatments. Estimated marginal means (EMMs) were calculated from an ANOVA model including treatment, nitrogen level, and block effects (n= 53). Pairwise differences between treatments were evaluated separately within each nitrogen level using Tukey's HSD test. The results below are from samples without added nitrogen (Nitrogen -). Data shown in Figure 2.

| *contrast* | *estimate* | *SE* | *df* | *t ratio* | *p value* |
|---|---|---|---|---|---|
| *L - LB* | -6.553 | 2.03 | 44 | -3.222 | 1.24E-02* |
| *L - LG* | -6.807 | 1.95 | 44 | -3.483 | 0.006* |
| *L - LGB* | -6.925 | 2.03 | 44 | -3.404 | 0.0075* |
| *LB – LGB* | -0.254 | 2.03 | 44 | -0.125 | 0.9993 |
| *LB – LGB* | -0.372 | 2.11 | 44 | -0.176 | 0.998 |
| *LG - LGB* | -0.118 | 2.03 | 44 | -0.058 | 0.9999 |

*Significant p<0.05

*Table S12:* Pairwise comparisons of the percentage of nitrogen from biological nitrogen fixation (BNF ) across treatments within nitrogen level. Estimated marginal means (EMMs) and standard errors were calculated from an ANOVA model including treatment, nitrogen level, and block effects (n= 53). Pairwise differences between treatments were evaluated separately within each nitrogen level using Tukey's HSD test.  Samples with added nitrogen (nitrogen +) are shown below. Data shown in Figure 2.

| *contrast* | *estimate* | *SE* | *df* | *t ratio* | *p value* |
|---|---|---|---|---|---|
| *L - LB* | -13.171 | 2.05 | 44 | -6.44 | <0.0001* |
| *L - LG* | -13.609 | 1.96 | 44 | -6.96 | <0.0001* |
| *L - LGB* | -21.515 | 2 | 44 | -10.757 | <0.0001* |
| *LB - LG* | -0.438 | 2.04 | 44 | -0.215 | 0.9964 |
| *LB - LGB* | -8.343 | 2.05 | 44 | -4.08 | 0.001* |
| *LG - LGB* | -7.905 | 1.99 | 44 | -3.976 | 0.0014* |

*Significant p<0.05

*Table S13:* ANOVA results on the main model for the nitrogen fixed per individual legume (grams). Model: N fixed per legume~ treatment* nitrogen + block. Data shown in Figure 2.

| *Term* | *df* | *Sum Sq* | *Mean Sq* | *F value* | *p value* |
|---|---|---|---|---|---|
| *Treatment* | 3 | 0.001323 | 0.000441 | 11.025 | 1.60E-05* |
| *Nitrogen* | 1 | 0.000762 | 0.000762 | 19.051 | 7.60E-05* |
| *Block* | 1 | 0.000595 | 0.000595 | 14.882 | 0.00037* |
| *Treatment: Nitrogen* | 3 | 0.000235 | 7.83E-05 | 1.956 | 0.13451 |
| *Residuals* | 44 | 0.001761 | 0.00004 | | |

*Significant p<0.05

*Table S14:* Pairwise comparisons of the percentage of nitrogen fixed per individual legume (grams) across treatments. Estimated marginal means (EMMs) were calculated from an ANOVA model including treatment, nitrogen level, and block effects (n= 53). Pairwise differences between treatments were evaluated separately within each nitrogen level using Tukey's HSD test. Samples without added nitrogen are shown below. Data shown in Figure 2.

| *contrast* | *estimate* | *SE* | *df* | *t ratio* | *p value* |
|---|---|---|---|---|---|
| *L - LB* | -0.00446 | 0.00352 | 44 | -1.268 | 0.5879 |
| *L - LG* | -0.0061 | 0.00338 | 44 | -1.805 | 0.2848 |
| *L - LGB* | -0.01806 | 0.00352 | 44 | -5.132 | <0.0001* |
| *LB - LG* | -0.00164 | 0.00352 | 44 | -0.466 | 0.9661 |
| *LB - LGB* | -0.0136 | 0.00365 | 44 | -3.724 | 0.003* |
| *LG - LGB* | -0.01196 | 0.00352 | 44 | -3.398 | 0.0076* |

*Significant p<0.05

*Table S15:* Pairwise comparisons of the percentage of nitrogen fixed per individual legume (grams) across treatments within nitrogen level. Estimated marginal means (EMMs) and standard errors were calculated from an ANOVA model including treatment, nitrogen level, and block effects (n= 53). Pairwise differences between treatments were evaluated separately within each nitrogen level using Tukey's HSD test. Samples with added nitrogen are shown below. Data shown in Figure 2.

| *contrast* | *estimate* | *SE* | *df* | *t ratio* | *p value* |
|---|---|---|---|---|---|
| *L - LB* | -0.01135 | 0.00354 | 44 | -3.208 | 0.0128* |
| *L - LG* | -0.00398 | 0.00338 | 44 | -1.176 | 0.6452 |
| *L - LGB* | -0.01317 | 0.00346 | 44 | -3.805 | 0.0024* |
| *LB - LG* | 0.00737 | 0.00353 | 44 | 2.09 | 0.1723 |
| *LB - LGB* | -0.00181 | 0.00354 | 44 | -0.512 | 0.9557 |
| *LG - LGB* | -0.00919 | 0.00344 | 44 | -2.671 | 0.0498* |

*Significant p<0.05

*Table S16:* Tukey-Sidak post-hoc test results for foxtail seed germination between treatments. The main model was a binomial model (germinating seeds: non-germinating seeds~ Treatment+ block), which had a significant effect of treatment (Type II Wald chi square test, p= 0.005). Results on the logit scale. Data shown in Figure S6.

| *contrast* | *estimate* | *SE* | *df* | *Z ratio* | *p value* |
|---|---|---|---|---|---|
| *L - G* | -0.0314 | 0.542 | Inf | -0.058 | 1 |
| *L - B* | 0.071 | 0.527 | Inf | 0.135 | 1 |
| *L - GB* | 0.3238 | 0.525 | Inf | 0.617 | 0.9963 |
| *L - LB* | 0.1945 | 0.542 | Inf | 0.359 | 0.9998 |
| *L - LG* | 0.4372 | 0.512 | Inf | 0.855 | 0.979 |
| *L - LGB* | 1.3682 | 0.446 | Inf | 3.071 | 0.0348* |
| *G - B* | 0.1025 | 0.525 | Inf | 0.195 | 1 |
| *G - GB* | 0.3552 | 0.526 | Inf | 0.676 | 0.9939 |
| *G - LB* | 0.226 | 0.542 | Inf | 0.417 | 0.9996 |
| *G - LG* | 0.4686 | 0.512 | Inf | 0.915 | 0.9704 |
| *G - LGB* | 1.3997 | 0.442 | Inf | 3.165 | 0.026* |
| *B - GB* | 0.2527 | 0.511 | Inf | 0.495 | 0.9989 |
| *B - LB* | 0.1235 | 0.528 | Inf | 0.234 | 1 |
| *B - LG* | 0.3662 | 0.497 | Inf | 0.737 | 0.9903 |
| *B - LGB* | 1.2972 | 0.42 | Inf | 3.088 | 0.0331* |
| *GB - LB* | -0.1292 | 0.526 | Inf | -0.246 | 1 |
| *GB - LG* | 0.1134 | 0.495 | Inf | 0.229 | 1 |
| *GB - LGB* | 1.0445 | 0.426 | Inf | 2.453 | 0.1767 |
| *LB - LG* | 0.2426 | 0.512 | Inf | 0.474 | 0.9992 |
| *LB - LGB* | 1.1737 | 0.446 | Inf | 2.63 | 0.1169 |
| *LG - LGB* | 0.931 | 0.409 | Inf | 2.276 | 0.2556 |

*Significant differences (p<0.05)

*Table S17:* Tukey-Sidak post-hoc test results for pigweed seed germination between treatments. The main model was a binomial model (germinating seeds: non-germinating seeds~ Treatment + block), which had a significant effect of treatment (Type II Wald chi square test, p < 0.001). Results on the logit scale. Data shown in Figure S6.

| *contrast* | *estimate* | *SE* | *df* | *z ratio* | *p value* |
|---|---|---|---|---|---|
| *L - G* | -0.1081 | 0.172 | Inf | -0.63 | 0.9959 |
| *L - B* | 0.7089 | 0.17 | Inf | 4.165 | 0.0006 |
| *L - GB* | -0.4297 | 0.187 | Inf | -2.294 | 0.2468 |
| *L - LB* | -0.6473 | 0.184 | Inf | -3.519 | 0.0079 |
| *L - LG* | -0.1267 | 0.18 | Inf | -0.703 | 0.9925 |
| *L - LGB* | 0.2116 | 0.18 | Inf | 1.177 | 0.903 |
| *G - B* | 0.817 | 0.168 | Inf | 4.85 | <0.0001* |
| *G - GB* | -0.3216 | 0.188 | Inf | -1.715 | 0.6057 |
| *G - LB* | -0.5391 | 0.184 | Inf | -2.934 | 0.052 |
| *G - LG* | -0.0186 | 0.18 | Inf | -0.103 | 1 |
| *G - LGB* | 0.3197 | 0.178 | Inf | 1.796 | 0.5505 |
| *B - GB* | -1.1387 | 0.186 | Inf | -6.114 | <0.0001* |
| *B - LB* | -1.3562 | 0.182 | Inf | -7.448 | <0.0001* |
| *B - LG* | -0.8356 | 0.179 | Inf | -4.668 | <0.0001* |
| *B - LGB* | -0.4973 | 0.175 | Inf | -2.844 | 0.0671 |
| *GB - LB* | -0.2175 | 0.199 | Inf | -1.095 | 0.9299 |
| *GB - LG* | 0.303 | 0.195 | Inf | 1.552 | 0.7132 |
| *GB - LGB* | 0.6414 | 0.195 | Inf | 3.288 | 0.0175 |
| *LB - LG* | 0.5206 | 0.192 | Inf | 2.711 | 0.0955 |
| *LB - LGB* | 0.8589 | 0.191 | Inf | 4.496 | 0.0001 |
| *LG - LGB* | 0.3383 | 0.188 | Inf | 1.798 | 0.5492 |

*Significant differences (p<0.05)

*Table S18:* Pairwise differences between expected and measured values for foxtail seed germination. The main model was binomial (germinating seeds: non-germinating seed ~ measurement type; expected - measured). Results on the logit scale. Data shown in Figure S6.

| *Treatment* | *contrast* | *df* | *Estimate* | *Std. Error* | *Z value* | *Pry(>\|z\|)* |
|---|---|---|---|---|---|---|
| *GB* | expected vs measured | 10 | 0.1885 | 0.5075 | -0.371 | 0.71 |
| *LB* | expected vs measured | 10 | 0.2029 | 0.5416 | -0.375 | 0.71 |
| *LG* | expected vs measured | 10 | 0.5933 | 0.5342 | -1.111 | 0.267 |
| *LGB* | expected vs measured | 10 | 1.599 | 0.4669 | -3.425 | 0.001* |

*Significant differences (p<0.05)

*Table S19:* Pairwise differences between expected and measured values for pigweed seed germination. The main model was a binomial model (germinating seeds: non-germinating seed ~ measurement type, expected - measured). Results on the logit scale. Data shown in Figure S6.

| *Treatment* | *contrast* | *df* | *Estimate* | *Std. Error* | *Z value* | *Pr(>\|z\|)* |
|---|---|---|---|---|---|---|
| *GB* | expected vs measured | 10 | -0.6448 | 0.1832 | -3.52 | 4.31E-04* |
| *LB* | expected vs measured | 11 | -0.9694 | 0.1805 | -5.37 | 7.89E-08* |
| *LG* | expected vs measured | 10 | -0.04917 | 0.17872 | -0.275 | 0.783 |
| *LGB* | expected vs measured | 10 | -0.05916 | 0.17444 | -0.339 | 0.734 |

*Significant differences (p<0.05)

*Table S20:* ANOVA results for the linear model of Shannon diversity in the total rhizosphere microbiome. Block was not significant, so it was removed from the model. Shannon diversity is calculated on rarefied reads with the R package Phyloseq. Model formula: Shannon diversity~ treatment* nitrogen. Results shown in Figure S2

| | *Df* | *Sum Sq* | *Mean Sq* | *F value* | *Pr(>F)* |
|---|---|---|---|---|---|
| *Treatment* | 6 | 0.8367 | 0.13945 | 5.85 | 5.67E-05* |
| *Nitrogen* | 1 | 0.1614 | 0.16136 | 6.769 | 0.0113* |
| *Treatment* Nitrogen* | 6 | 0.0872 | 0.01454 | 0.61 | 0.7216 |
| *Residuals* | 69 | 1.6448 | 0.02384 | | |

*Significant differences (p<0.05)

*Table S21:* ANOVA results of the linear model for ASV richness of the total rhizosphere microbiome. Block was not significant, so it was removed from the model. ASV richness is calculated on rarefied reads. Model formula: ASV richness ~ treatment* nitrogen. Results shown in Figure S2.

| | *Df* | *Sum Sq* | *Mean Sq* | *F value* | *Pr(>F)* |
|---|---|---|---|---|---|
| *Treatment* | 6 | 6068684 | 1011447 | 3.17E+00 | 0.00832* |
| *Nitrogen* | 1 | 1626724 | 1626724 | 5.1 | 0.02709* |
| *Treatment* nitrogen* | 6 | 1735024 | 289171 | 0.907 | 0.4954 |
| *Residuals* | 69 | 22007168 | 318944 | | |

*Significant differences (p<0.05)

*Table S22:* ANOVA results of the linear model for Pilou's Evenness (J) of the total rhizosphere microbiome. Block was not significant, so it was removed from the model. Evenness is calculated on rarefied reads as J = H'/ln(S), where H' is the Shannon diversity and S is the number of species. Model formula: evenness~ treatment* nitrogen. Results shown in Figure S2.

| | *Df* | *Sum Sq* | *Mean Sq* | *F value* | *Pr(>F)* |
|---|---|---|---|---|---|
| *Treatment* | 6 | 0.002023 | 0.0003372 | 3.697 | 0.00304* |
| *Nitrogen* | 1 | 0.000156 | 0.0001562 | 1.713 | 0.19492 |
| *Treatment* nitrogen* | 6 | 0.000534 | 0.0000891 | 0.977 | 0.4477 |
| *Residuals* | 69 | 0.006292 | 0.0000912 | | |

*Significant differences (p<0.05)

*Table S23:* Tukey post hoc comparisons Shannon diversity of total rhizosphere microbiome. Model formula: Shannon diversity ~ treatment. Results shown in Figure S2.

| *contrast* | *estimate* | *SE* | *df* | *t ratio* | *p.value* |
|---|---|---|---|---|---|
| *L - G* | -0.26271 | 0.0659 | 76 | -3.987 | 0.0028* |
| *L - B* | -0.33934 | 0.0659 | 76 | -5.15 | <0.0001* |
| *L - GB* | -0.27108 | 0.0659 | 76 | -4.114 | 0.0018* |
| *L - LB* | -0.24471 | 0.0659 | 76 | -3.714 | 0.0068* |
| *L - LG* | -0.2912 | 0.0659 | 76 | -4.42 | 0.0006* |
| *L - LGB* | -0.17942 | 0.0659 | 76 | -2.723 | 0.1064 |
| *G - B* | -0.07663 | 0.0644 | 76 | -1.189 | 0.8961 |
| *G - GB* | -0.00837 | 0.0644 | 76 | -0.13 | 1 |
| *G - LB* | 0.018 | 0.0644 | 76 | 0.279 | 1 |
| *G - LG* | -0.02849 | 0.0644 | 76 | -0.442 | 0.9994 |
| *G - LGB* | 0.08328 | 0.0644 | 76 | 1.292 | 0.8533 |
| *B - GB* | 0.06826 | 0.0644 | 76 | 1.059 | 0.9378 |
| *B - LB* | 0.09463 | 0.0644 | 76 | 1.469 | 0.7622 |
| *B - LG* | 0.04814 | 0.0644 | 76 | 0.747 | 0.989 |
| *B - LGB* | 0.15991 | 0.0644 | 76 | 2.482 | 0.1807 |
| *GB - LB* | 0.02637 | 0.0644 | 76 | 0.409 | 0.9996 |
| *GB - LG* | -0.02012 | 0.0644 | 76 | -0.312 | 0.9999 |
| *GB - LGB* | 0.09165 | 0.0644 | 76 | 1.422 | 0.7881 |
| *LB - LG* | -0.04649 | 0.0644 | 76 | -0.722 | 0.9909 |
| *LB - LGB* | 0.06528 | 0.0644 | 76 | 1.013 | 0.9495 |
| *LG - LGB* | 0.11178 | 0.0644 | 76 | 1.735 | 0.5954 |

*Significant differences (p<0.05)

*Table S24:* Tukey post hoc comparisons for ASV richness of the total rhizosphere microbiome. Model formula: ASV richness~ treatment. Results shown in Figure S2.

| *contrast* | *estimate* | *SE* | *df* | *t.ratio* | *p.value* |
|---|---|---|---|---|---|
| *L - G* | -587.3 | 241 | 76 | -2.435 | 0.1986 |
| *L - B* | -798.7 | 241 | 76 | -3.312 | 0.023* |
| *L - GB* | -775.8 | 241 | 76 | -3.217 | 0.0301 |
| *L - LB* | -729.5 | 241 | 76 | -3.025 | 0.0505 |
| *L - LG* | -815 | 241 | 76 | -3.38 | 0.0189 |
| *L - LGB* | -393.8 | 241 | 76 | -1.633 | 0.6618 |
| *G - B* | -211.4 | 236 | 76 | -0.896 | 0.9721 |
| *G - GB* | -188.5 | 236 | 76 | -0.799 | 0.9844 |
| *G - LB* | -142.2 | 236 | 76 | -0.603 | 0.9965 |
| *G - LG* | -227.8 | 236 | 76 | -0.966 | 0.9598 |
| *G - LGB* | 193.5 | 236 | 76 | 0.82 | 0.9821 |
| *B - GB* | 22.9 | 236 | 76 | 0.097 | 1 |
| *B - LB* | 69.2 | 236 | 76 | 0.293 | 0.9999 |
| *B - LG* | -16.3 | 236 | 76 | -0.069 | 1 |
| *B - LGB* | 404.9 | 236 | 76 | 1.717 | 0.6072 |
| *GB - LB* | 46.2 | 236 | 76 | 0.196 | 1 |
| *GB - LG* | -39.2 | 236 | 76 | -0.166 | 1 |
| *GB - LGB* | 382 | 236 | 76 | 1.62 | 0.6703 |
| *LB - LG* | -85.5 | 236 | 76 | -0.362 | 0.9998 |
| *LB - LGB* | 335.8 | 236 | 76 | 1.423 | 0.7875 |
| *LG - LGB* | 421.2 | 236 | 76 | 1.786 | 0.5615 |

*Table S25:* Tukey post hoc comparisons for Pilou's evenness (J) of the total rhizosphere microbiome. Model formula: evenness~ treatment. Results shown in Figure S2.

| *contrast* | *estimate* | *SE* | *df* | *t.ratio* | *p.value* |
|---|---|---|---|---|---|
| *L - G* | -0.014285 | 0.004 | 76 | -3.57 | 0.0107* |
| *L - B* | -0.016934 | 0.004 | 76 | -4.232 | 0.0012* |
| *L - GB* | -0.008856 | 0.004 | 76 | -2.214 | 0.3007 |
| *L - LB* | -0.007224 | 0.004 | 76 | -1.805 | 0.5486 |
| *L - LG* | -0.010787 | 0.004 | 76 | -2.696 | 0.1133 |
| *L - LGB* | -0.008714 | 0.004 | 76 | -2.178 | 0.3196 |
| *G - B* | -0.002649 | 0.00391 | 76 | -0.677 | 0.9935 |
| *G - GB* | 0.005428 | 0.00391 | 76 | 1.387 | 0.8069 |
| *G - LB* | 0.007061 | 0.00391 | 76 | 1.804 | 0.5493 |
| *G - LG* | 0.003498 | 0.00391 | 76 | 0.894 | 0.9724 |
| *G - LGB* | 0.00557 | 0.00391 | 76 | 1.424 | 0.7875 |
| *B - GB* | 0.008077 | 0.00391 | 76 | 2.064 | 0.3843 |
| *B - LB* | 0.00971 | 0.00391 | 76 | 2.481 | 0.1808 |
| *B - LG* | 0.006147 | 0.00391 | 76 | 1.571 | 0.701 |
| *B - LGB* | 0.008219 | 0.00391 | 76 | 2.1 | 0.363 |
| *GB - LB* | 0.001633 | 0.00391 | 76 | 0.417 | 0.9996 |
| *GB - LG* | -0.001931 | 0.00391 | 76 | -0.493 | 0.9989 |
| *GB - LGB* | 0.000142 | 0.00391 | 76 | 0.036 | 1 |
| *LB - LG* | -0.003563 | 0.00391 | 76 | -0.911 | 0.9698 |
| *LB - LGB* | -0.001491 | 0.00391 | 76 | -0.381 | 0.9997 |
| *LG - LGB* | 0.002073 | 0.00391 | 76 | 0.53 | 0.9983 |

*Significant differences ($p<0.05$)

*Table S26:* Permutation test on the Constrained Analysis of Principal Coordinates (CAP) model of Bray-Curtis dissimilarities, for the total rhizosphere community. Model formula: distance matrix ~ treatment* nitrogen + block. Data shown in Figure 3.

| *Term* | *df* | *Sum of Sqs* | *F value* | *p value* |
|---|---|---|---|---|
| *Treatment* | 6 | 2.7401 | 3.3524 | 0.001* |
| *Nitrogen* | 1 | 0.4135 | 3.0354 | 0.001* |
| *Block* | 1 | 0.4123 | 3.0268 | 0.001* |
| *Treatment* Nitrogen* | 6 | 0.9618 | 1.1767 | 0.052 |
| *Residual* | 68 | 9.2634 | | |

*Significant differences (p<0.05)

*Table S27:* Pairwise permutation tests for the CAP analysis of Bray-Curtis dissimilarities for the total rhizosphere community. Data for samples without exogenous nitrogen applied. P-values are corrected for false discovery rate (FDR) for multiple testing. Model formula: distance matrix ~ treatment + block. Data shown in Figure 3.

| *contrast* | *Df* | *Sum of sqs.* | *F value* | *p value* |
|---|---|---|---|---|
| *L - G* | 1 | 0.56825 | 3.9667 | 0.003* |
| *L - B* | 1 | 0.63744 | 4.9467 | 0.003* |
| *L - GB* | 1 | 0.58999 | 4.2179 | 0.003* |
| *L - LB* | 1 | 0.18047 | 1.3396 | 0.0968 |
| *L - LG* | 1 | 0.2792 | 2.1453 | 0.003* |
| *L - LGB* | 1 | 0.24316 | 1.8136 | 0.0035* |
| *G - B* | 1 | 0.19623 | 1.3228 | 0.0035* |
| *G - GB* | 1 | 0.14955 | 0.9384 | 0.19 |
| *G - LB* | 1 | 0.34008 | 2.2053 | 0.0048* |
| *G - LG* | 1 | 0.24531 | 1.6394 | 0.007* |
| *G - LGB* | 1 | 0.28261 | 1.8404 | 0.0035* |
| *B - GB* | 1 | 0.18601 | 1.2831 | 0.0105* |
| *B - LB* | 1 | 0.37206 | 2.6611 | 0.0035* |
| *B - LG* | 1 | 0.34047 | 2.5175 | 0.003* |
| *B - LGB* | 1 | 0.37675 | 2.7071 | 0.003* |
| *GB - LB* | 1 | 0.36978 | 2.4516 | 0.007* |
| *GB - LG* | 1 | 0.28881 | 1.9746 | 0.0035* |
| *GB - LGB* | 1 | 0.33301 | 2.2173 | 0.003* |
| *LB - LG* | 1 | 0.15289 | 1.0836 | 0.1802 |
| *LB - LGB* | 1 | 0.13646 | 0.9409 | 0.1869 |
| *LG - LGB* | 1 | 0.14669 | 1.0443 | 0.0383* |

*Significant differences (p<0.05)

*Table S28:* Pairwise permutation tests for the CAP model of Bray-Curtis dissimilarities for the total rhizosphere community. P-values are FDR (false discovery rate) corrected. Data for nitrogen-fertilized samples only. (Model: distance matrix ~ treatment + block). Data shown in Figure 3.

| *contrast* | *Df* | *Sum of Sqs* | *F Value* | *p value* |
|---|---|---|---|---|
| *L - G* | 1 | 0.5148 | 3.586 | 0.0013* |
| *L - B* | 1 | 0.43793 | 3.0191 | 0.0013* |
| *L - GB* | 1 | 0.51008 | 3.352 | 0.0013* |
| *L - LB* | 1 | 0.17257 | 1.2508 | 0.0013* |
| *L - LG* | 1 | 0.27104 | 1.8721 | 0.0022* |
| *L - LGB* | 1 | 0.28231 | 1.918 | 0.0013* |
| *G - B* | 1 | 0.34626 | 2.6094 | 0.0013* |
| *G - GB* | 1 | 0.16409 | 1.1796 | 0.0032* |
| *G - LB* | 1 | 0.49849 | 3.9462 | 0.0013* |
| *G - LG* | 1 | 0.31241 | 2.3588 | 0.0013* |
| *G - LGB* | 1 | 0.3078 | 2.2865 | 0.0013* |
| *B - GB* | 1 | 0.33945 | 2.4169 | 0.0013* |
| *B - LB* | 1 | 0.32509 | 2.5465 | 0.0013* |
| *B - LG* | 1 | 0.32694 | 2.4436 | 0.0022* |
| *B - LGB* | 1 | 0.2529 | 1.8601 | 0.0013* |
| *GB - LB* | 1 | 0.44693 | 3.3335 | 0.0013* |
| *GB - LG* | 1 | 0.26467 | 1.8878 | 0.0013* |
| *GB - LGB* | 1 | 0.29373 | 2.0632 | 0.0013* |
| *LB - LG* | 1 | 0.18105 | 1.421 | 0.0013* |
| *LB - LGB* | 1 | 0.17876 | 1.3795 | 0.0022* |
| *LG - LGB* | 1 | 0.13306 | 0.9805 | 0.042* |

*Significant differences ($p<0.05$)

*Table S29:* ANOVA-like permutation test results of the CAP analysis on the effect of each functional group in the total microbial community for unfertilized samples only. Model: distance matrix ~ grass * brassicae * legume + block).

| | *Df* | *Sum of Sqs* | *F* | *Pr(>F)* |
|---|---|---|---|---|
| *Legume* | 1 | 0.909 | 6.4917 | 0.001* |
| *Grass* | 1 | 0.2541 | 1.8149 | 0.015* |
| *Brassicae* | 1 | 0.1659 | 1.1847 | 0.162 |
| *Block* | 1 | 0.2554 | 1.8242 | 0.011* |
| *Legume*Grass* | 1 | 0.1761 | 1.2578 | 0.138 |
| *Legume*Brassicae* | 1 | 0.1537 | 1.0974 | 0.255 |
| *Grass*Brassicae* | 1 | 0.1734 | 1.2383 | 0.128 |
| *Residual* | 34 | 4.7606 | | |

*Significant differences ($p<0.05$)

*Table S30:* ANOVA-like permutation test results of the distance-based redundancy analysis on the effect of each functional group in the nitrogen-fertilized total microbial community (Nitrogen +). Model: distance matrix ~ grass * brassicae * legume + block).

| | *Df* | *Sum of Sqs* | *F* | *Pr(>F)* |
|---|---|---|---|---|
| *Legume* | 1 | 0.6959 | 5.2486 | 0.001* |
| *Grass* | 1 | 0.4301 | 3.2443 | 0.001* |
| *Brassicae* | 1 | 0.1989 | 1.5002 | 0.036* |
| *Block* | 1 | 0.2827 | 2.1326 | 0.003* |
| *Legume*Grass* | 1 | 0.2328 | 1.7556 | 0.007* |
| *Legume*Brassicae* | 1 | 0.1626 | 1.2264 | 0.132 |
| *Grass*Brassicae* | 1 | 0.1487 | 1.1215 | 0.208 |
| *Residual* | 33 | 4.3751 | | |

*Significant differences ($p<0.05$)

*Table S31:* ANOVA between expected and measured values of centered log-transformed microbial composition scaled between monocultures on the total nitrogen fertilized microbial community. munity. Rep included to account for the paired nature of samples. Model: distance ~ measured vs. expected + rep).

| *Mixture* | *Df (test, residual)* | *Sum of Sqs* | *Mean Sq.* | *F* | *Pr(>F)* |
|---|---|---|---|---|---|
| *Grass Brassica* | 1, 4 | 0.02056 | 0.02056 | 5.232 | 0.0841 |
| *Legume Brassica* | 1, 4 | 0.08179 | 0.08179 | 3.177 | 0.149 |
| *Legume Grass* | 1, 4 | 0.01802 | 0.01802 | 0.421 | 0.552 |
| *Three species mix- GB index* | 1,7 | 0.0349 | 0.0349 | 9.269 | 0.0187* |
| *Three species mix- LB index* | 1,7 | 0.01632 | 0.016317 | 2.357 | 0.169 |
| *Three species mix- LG index* | 1,7 | 0.00075 | 0.000749 | 0.089 | 0.774 |

*Significant differences (p<0.05)

*Table S32:* Model of the effect of legume presence/absence on the number of active cells per gram of rhizosphere. The main model was a negative binomial (active cells/g~ legume + block).

| | *Estimate* | *Std. Error* | *z value* | *Pr(>\|z\|)* |
|---|---|---|---|---|
| *(Intercept)* | 5.8534 | 0.3007 | 19.467 | < 2e-16* |
| *Legume* | 1.5906 | 0.2484 | 6.404 | 1.51E-10* |
| *block* | -0.1561 | 0.1024 | -1.525 | 0.127 |

*Significant differences (p<0.05)

*Table S33:* Model of the effect of Legume presence/absence on percent active cells in a binomial model (percent active ~ legume + block ).

| | *Estimate* | *Std. Error* | *z value* | *Pr(>\|z\|)* |
|---|---|---|---|---|
| *(Intercept)* | -2.86955 | 0.012468 | -230.144 | < 2e-16* |
| *Legume* | 0.062586 | 0.010705 | 5.846 | 5.03E-09* |
| *block* | 0.05687 | 0.004374 | 13.001 | < 2e-16* |

*Significant differences (p<0.05)

*Table S34:* Tukey comparison for the number of active cells per gram of rhizosphere. The main model was a negative binomial model (formula: percent active cells~ treatment + block). Data shown in Figure 4.

| *contrast* | *estimate* | *SE* | *df* | *Z ratio* | *p value* |
|---|---|---|---|---|---|
| *L - G* | 2.0611 | 0.419 | Inf | 4.923 | <0.0001* |
| *L - B* | 1.66056 | 0.467 | Inf | 3.559 | 0.0068* |
| *L - GB* | 1.66351 | 0.437 | Inf | 3.804 | 0.0027* |
| *L - LB* | 0.27213 | 0.437 | Inf | 0.622 | 0.9961 |
| *L - LG* | 0.27522 | 0.437 | Inf | 0.63 | 0.9958 |
| *L - LGB* | 0.47149 | 0.422 | Inf | 1.117 | 0.923 |
| *G - B* | -0.40053 | 0.47 | Inf | -0.853 | 0.9792 |
| *G - GB* | -0.39759 | 0.439 | Inf | -0.905 | 0.9719 |
| *G - LB* | -1.78897 | 0.438 | Inf | -4.087 | 0.0009* |
| *G - LG* | -1.78588 | 0.439 | Inf | -4.071 | 0.0009* |
| *G - LGB* | -1.58961 | 0.419 | Inf | -3.796 | 0.0028* |
| *B - GB* | 0.00295 | 0.485 | Inf | 0.006 | 1 |
| *B - LB* | -1.38843 | 0.486 | Inf | -2.856 | 0.0649 |
| *B - LG* | -1.38535 | 0.485 | Inf | -2.857 | 0.0647 |
| *B - LGB* | -1.18907 | 0.474 | Inf | -2.509 | 0.156 |
| *GB - LB* | -1.39138 | 0.457 | Inf | -3.043 | 0.0378* |
| *GB - LG* | -1.38829 | 0.457 | Inf | -3.04 | 0.0382* |
| *GB - LGB* | -1.19202 | 0.442 | Inf | -2.696 | 0.0992 |
| *LB - LG* | 0.00309 | 0.457 | Inf | 0.007 | 1 |
| *LB - LGB* | 0.19936 | 0.439 | Inf | 0.454 | 0.9993 |
| *LG - LGB* | 0.19627 | 0.442 | Inf | 0.445 | 0.9994 |

*Significant differences (p<0.05)

*Table S35:* Pairwise differences between expected and measured values for the number of active cells per gram of rhizosphere. The main model was a negative binomial model (number of active cells/g ~ measurement type, expected vs. measured +rep ). Data shown in Figure 4.

| *Treatment* | *contrast* | *Df (term, residual)* | *Estimate* | *Std.Error* | *zvalue* | *Pr(>\|z\|)* |
|---|---|---|---|---|---|---|
| *GB* | expected vs measured | 1,7 | 0.572 | 0.316 | 1.81 | 0.0709 |
| *LB* | expected vs measured | 1,7 | 0.663 | 0.489 | 1.35 | 0.176 |
| *LG* | expected vs measured | 1,9 | 0.174 | 0.418 | 0.417 | 0.677 |
| *LGB* | expected vs measured | 1,8 | 0.488 | 0.482 | 1.01 | 0.311 |

*Significant differences (p<0.05)

*Table S36:* Tukey comparison for the percentage of active cells. The main model was binomial (formula: active cells:total cells ~ Treatment + block). Data shown in Figure 4.

| *contrast* | *estimate* | *SE* | *df* | *z.ratio* | *p.value* |
|---|---|---|---|---|---|
| *L - G* | 0.05086 | 0.0203 | Inf | 2.51E+00 | 0.1569 |
| *L - B* | -0.07605 | 0.0178 | Inf | -4.272 | 0.0004* |
| *L - GB* | 0.00103 | 0.0185 | Inf | 5.60E-02 | 1 |
| *L - LB* | -0.01261 | 0.0173 | Inf | -0.73 | 0.9907 |
| *L - LG* | -0.26767 | 0.0157 | Inf | -17.053 | <0.0001* |
| *L - LGB* | -0.043 | 0.0168 | Inf | -2.565 | 0.1368 |
| *G - B* | -0.12692 | 0.0224 | Inf | -5.656 | <0.0001* |
| *G - GB* | -0.04983 | 0.0232 | Inf | -2.15 | 0.3232 |
| *G - LB* | -0.06348 | 0.0222 | Inf | -2.864 | 0.0635 |
| *G - LG* | -0.31854 | 0.0208 | Inf | -15.325 | <0.0001* |
| *G - LGB* | -0.09387 | 0.0215 | Inf | -4.36 | 0.0003* |
| *B - GB* | 0.07709 | 0.021 | Inf | 3.679 | 0.0044* |
| *B - LB* | 0.06344 | 0.0199 | Inf | 3.194 | 0.0237* |
| *B - LG* | -0.19162 | 0.0184 | Inf | -10.419 | <0.0001* |
| *B - LGB* | 0.03305 | 0.0193 | Inf | 1.716 | 0.6051 |
| *GB - LB* | -0.01365 | 0.0201 | Inf | -0.678 | 0.9938 |
| *GB - LG* | -0.26871 | 0.0192 | Inf | -13.96 | <0.0001* |
| *GB - LGB* | -0.04404 | 0.0203 | Inf | -2.169 | 0.3127 |
| *LB - LG* | -0.25506 | 0.018 | Inf | -14.143 | <0.0001* |
| *LB - LGB* | -0.03039 | 0.0191 | Inf | -1.592 | 0.6873 |
| *LG - LGB* | 0.22467 | 0.0173 | Inf | 12.997 | <0.0001* |

*Significant differences (p<0.05)

*Table S37:* Pairwise differences between expected and measured values for percent microbial activity. The main model was binomial (formula: percent active cells ~ measurement type, expected vs. measured + rep). Results on the logit scale. Data shown in Figure 4.

| *Treatment* | *contrast* | *Df (term, residual)* | *Estimate* | *Std.Error* | *zvalue* | *Pr(>\|z\|)* |
|---|---|---|---|---|---|---|
| *GB* | expected vs measured | 1,6 | 0.19053 | 0.27133 | 0.702 | 0.483 |
| *LB* | expected vs measured | 1,6 | -0.33862 | 0.29261 | -1.157 | 0.247 |
| *LG* | expected vs measured | 1,8 | -0.59665 | 0.23208 | -2.571 | 0.0101* |
| *LGB* | expected vs measured | 1,8 | -0.01324 | 0.28309 | -0.047 | 0.9627 |

*Significant differences (p<0.05)

*Table S38:* Overall model for active vs inactive rhizosphere microbiomes. Permutation test on CAP ordination of Bray-Curtis dissimilarities. (Model: Distance matrix~ Treatment * Fraction + block). Data shown in Figure 4.

| | *df* | *sum of sqs* | *F* | *p value* |
|---|---|---|---|---|
| *Treatment* | 6 | 2.719 | 1.7706 | 0.001* |
| *Fraction* | 1 | 1.0959 | 4.282 | 0.001* |
| *Block* | 1 | 0.7564 | 2.9556 | 0.001* |
| *Treatment*Fraction* | 6 | 1.5223 | 0.9914 | 0.532 |
| *Residual* | 52 | 13.3085 | | |

*Significant differences (p<0.05

*Table S39:* Pairwise permutation test for CAP ordination of the inactive rhizosphere microbial community (distance matrix ~ treatment + conditioned(block).

| *contrast* | *Df* | *SumOfSqs* | *F* | *Pr(>F)* | *Fdr adjusted p values* |
|---|---|---|---|---|---|
| *L - G* | 1 | 0.45089 | 1.6766 | 0.031 | 0.1085 |
| *L - B* | 1 | 0.42653 | 1.3991 | 0.117 | 0.2424545 |
| *L - GB* | 1 | 0.34103 | 1.2366 | 0.185 | 0.2988462 |
| *L - LB* | 1 | 0.29994 | 1.1193 | 0.303 | 0.4242 |
| *L - LG* | 1 | 0.36685 | 1.3119 | 0.083 | 0.203 |
| *L - LGB* | 1 | 0.26203 | 0.8255 | 0.674 | 0.7077 |
| *G - B* | 1 | 0.45288 | 1.635 | 0.008 | 0.0672 |
| *G - GB* | 1 | 0.19664 | 0.8404 | 0.966 | 0.966 |
| *G - LB* | 1 | 0.5938 | 2.5097 | 0.011 | 0.0672 |
| *G - LG* | 1 | 0.40495 | 1.6311 | 0.012 | 0.0672 |
| *G - LGB* | 1 | 0.37554 | 1.3336 | 0.087 | 0.203 |
| *B - GB* | 1 | 0.29573 | 1.0371 | 0.419 | 0.5106316 |
| *B - LB* | 1 | 0.52308 | 1.8944 | 0.042 | 0.126 |
| *B - LG* | 1 | 0.33253 | 1.1607 | 0.166 | 0.2905 |
| *B - LGB* | 1 | 0.32129 | 1.0027 | 0.458 | 0.5106316 |
| *GB - LB* | 1 | 0.39575 | 1.7009 | 0.016 | 0.0672 |
| *GB - LG* | 1 | 0.30666 | 1.2354 | 0.127 | 0.2424545 |
| *GB - LGB* | 1 | 0.28338 | 0.961 | 0.462 | 0.5106316 |
| *LB - LG* | 1 | 0.4679 | 1.892 | 0.015 | 0.0672 |
| *LB - LGB* | 1 | 0.28377 | 1.0117 | 0.423 | 0.5106316 |
| *LG - LGB* | 1 | 0.33874 | 1.1529 | 0.232 | 0.348 |

*Significant differences ($p<0.05$)

*Table S40:* Pairwise permutation test for CAP ordination model for the active rhizosphere microbial community (distance matrix ~ Treatment + conditioned(block).

| *contrast* | *Df* | *SumOfSqs* | *F* | *Pr(>F)* | *Fdr adjusted p value* |
|---|---|---|---|---|---|
| *L - G* | 1 | 0.47882 | 2.089 | 0.012 | 0.042* |
| *L - B* | 1 | 0.46243 | 1.864 | 0.025 | 0.0735 |
| *L - GB* | 1 | 0.35043 | 1.3182 | 0.147 | 0.075 |
| *L - LB* | 1 | 0.27396 | 1.214 | 0.212 | 0.274909 |
| *L - LG* | 1 | 0.35998 | 1.4213 | 0.117 | 0.274909 |
| *L - LGB* | 1 | 0.27503 | 1.1908 | 0.3 | 0.4335 |
| *G - B* | 1 | 0.50991 | 2.0464 | 0.013 | 0.042* |
| *G - GB* | 1 | 0.25825 | 0.9744 | 0.439 | 0.4335 |
| *G - LB* | 1 | 0.48221 | 2.0993 | 0.006 | 0.042* |
| *G - LG* | 1 | 0.41763 | 1.6456 | 0.037 | 0.042* |
| *G - LGB* | 1 | 0.36596 | 1.5585 | 0.042 | 0.08925 |
| *B - GB* | 1 | 0.25722 | 0.9 | 0.637 | 0.496125 |
| *B - LB* | 1 | 0.37087 | 1.4756 | 0.017 | 0.042* |
| *B - LG* | 1 | 0.26034 | 0.9442 | 0.577 | 0.274909 |
| *B - LGB* | 1 | 0.25472 | 0.9763 | 0.378 | 0.639947 |
| *GB - LB* | 1 | 0.31764 | 1.1725 | 0.214 | 0.308 |
| *GB - LG* | 1 | 0.22225 | 0.7653 | 0.807 | 0.639947 |
| *GB - LGB* | 1 | 0.21766 | 0.7721 | 0.887 | 0.91 |
| *LB - LG* | 1 | 0.2487 | 0.9667 | 0.557 | 0.483 |
| *LB - LGB* | 1 | 0.23576 | 1.0158 | 0.409 | 0.541059 |
| *LG - LGB* | 1 | 0.20836 | 0.778 | 0.901 | 0.91 |

*Significant differences ($p<0.05$)

*Table S41:* Permutation test on CAP ordination of Bray-Curtis dissimilarities of the effect of each functional group in the active community (distance matrix ~ grass * brassicae * legume + condition(block)). Model terms are the binary variables of the present/absence of a plant functional group in a plant mixture.

| *Term* | *Df* | *SumOfSqs* | *F* | *Pr(>F)* |
|---|---|---|---|---|
| *Legume* | 1 | 0.5194 | 2.1595 | 0.002* |
| *Grass* | 1 | 0.3095 | 1.2867 | 0.138 |
| *Brassicae* | 1 | 0.3168 | 1.3171 | 0.105 |
| *Legume* Grass* | 1 | 0.3796 | 1.5781 | 0.034* |
| *Legume* Brassicae* | 1 | 0.2398 | 0.997 | 0.44 |
| *Grass* Brassicae* | 1 | 0.2538 | 1.0553 | 0.367 |
| *Residual* | 26 | 6.2533 | | |

*Significant differences (p<0.05)

*Table S42:* ANOVA between expected and measured values of centered log-transformed microbial composition for scaled between monocultures on the active microbial community. Pots were nitrogen-fertilized. Rep included to account for the paired nature of samples. Model formula is distance ~ measured vs. expected + rep.

| *Mixture* | *Df (test, residual)* | *Sum of Sqs* | *Mean Sq.* | *F* | *Pr(>F)* |
|---|---|---|---|---|---|
| *Grass Brassica* | 1,7 | 0.0681 | 0.0681 | 4.550 | 0.070 |
| *Legume Brassica* | 1,5 | 0.0335 | 0.0335 | 2.337 | 0.187 |
| *Legume Grass* | 1,7 | 0.0166 | 0.0166 | 0.291 | 0.606 |
| *Three species mix- GB index* | 1,4 | 0.02634 | 0.02634 | 4.955 | 0.09 |
| *Three species mix- LB index* | 1,4 | 0.02152 | 0.02152 | 0.896 | 0.397 |
| *Three species mix- LG index* | 1,4 | 0.00045 | 0.000454 | 0.021 | 0.891 |

*Significant differences (p<0.05)

*Table S43:* Differentially abundant taxa in the total nitrogen-fertilized community. First, twenty ASVs with the highest species scores on CAP 1 and CAP 2 from the CAP ordination of Bray-Curtis dissimilarities (distance matrix~ Treatment +block) were selected. These ASVs were aggregated to the lowest taxonomic level. Then DESeq2 was run on these key taxa (rarefied reads ~ legume * grass * grassica). Model terms are the binary variables of the present/absence of a plant functional group. The base mean (rarefied reads) and log fold change are calculated from the plant-free soil pots. Only significant taxa are shown (p adj < 0.01).

| Model term | Lowest taxonomic ID | Base Mean | log2 Fold Change | lfcSE | Wald test stat | P value | P adj |
|---|---|---|---|---|---|---|---|
| intercept | Arthrobacter sp. | 195.2 | -2.9 | 0.7 | -4.1 | 4.7E-05 | 3.7E-04 |
| L | Arthrobacter sp. | 195.2 | -3.9 | 0.5 | -8.6 | 8.2E-18 | 3.3E-17 |
| B | Arthrobacter sp. | 195.2 | -3.8 | 0.4 | -8.4 | 3.7E-17 | 3.0E-16 |
| G | Arthrobacter sp. | 195.2 | -3.6 | 0.4 | -8.0 | 1.7E-15 | 9.3E-15 |
| G*B | Arthrobacter sp. | 195.2 | 3.2 | 0.6 | 5.8 | 6.3E-09 | 5.1E-08 |
| L*B | Arthrobacter sp. | 195.2 | 3.7 | 0.6 | 6.7 | 2.7E-11 | 2.1E-10 |
| L*G | Arthrobacter sp. | 195.2 | 3.5 | 0.6 | 6.3 | 3.5E-10 | 1.4E-09 |
| L*G*B | Arthrobacter sp. | 195.2 | -2.9 | 0.7 | -4.1 | 4.7E-05 | 3.7E-04 |
| L | Caulobacter sp. | 169.4 | 4.1 | 0.9 | 4.3 | 1.8E-05 | 2.9E-05 |
| B | Caulobacter sp. | 169.4 | 3.3 | 0.9 | 3.5 | 5.3E-04 | 1.1E-03 |
| G | Caulobacter sp. | 169.4 | 3.4 | 0.9 | 3.6 | 2.8E-04 | 4.5E-04 |
| L*B | Caulobacter sp. | 169.4 | -3.0 | 1.0 | -2.9 | 3.5E-03 | 5.6E-03 |
| L*G | Caulobacter sp. | 169.4 | -3.6 | 1.0 | -3.4 | 5.7E-04 | 1.1E-03 |
| L | Limnocylindria | 447.7 | -3.4 | 0.4 | -7.9 | 2.5E-15 | 6.7E-15 |
| B | Limnocylindria | 447.7 | -2.3 | 0.4 | -5.5 | 4.5E-08 | 1.2E-07 |
| G | Limnocylindria | 447.7 | -2.7 | 0.4 | -6.4 | 1.7E-10 | 4.5E-10 |
| G*B | Limnocylindria | 447.7 | 2.0 | 0.5 | 3.9 | 7.8E-05 | 2.5E-04 |
| L*B | Limnocylindria | 447.7 | 2.2 | 0.5 | 4.1 | 3.5E-05 | 9.4E-05 |
| L*G | Limnocylindria | 447.7 | 2.8 | 0.5 | 5.3 | 1.3E-07 | 3.5E-07 |
| L | Nitrosocosmicus sp. | 1282.5 | -4.5 | 0.5 | -9.8 | 1.1E-22 | 8.5E-22 |
| B | Nitrosocosmicus sp. | 1282.5 | -2.8 | 0.5 | -6.3 | 3.0E-10 | 1.2E-09 |
| G | Nitrosocosmicus sp. | 1282.5 | -3.6 | 0.5 | -7.9 | 2.3E-15 | 9.3E-15 |
| G*B | Nitrosocosmicus sp. | 1282.5 | 2.2 | 0.6 | 3.9 | 9.5E-05 | 2.5E-04 |
| L*B | Nitrosocosmicus sp. | 1282.5 | 3.2 | 0.6 | 5.6 | 1.8E-08 | 7.3E-08 |
| L*G | Nitrosocosmicus sp. | 1282.5 | 3.7 | 0.6 | 6.5 | 7.4E-11 | 5.9E-10 |
| G | Polaromonas sp. | 54.4 | 4.5 | 1.4 | 3.3 | 9.9E-04 | 1.3E-03 |
| L | Rhizobium sp. | 617.5 | 3.9 | 0.8 | 4.6 | 3.3E-06 | 6.6E-06 |
| L*B | Rhizobium sp. | 617.5 | -3.1 | 1.0 | -3.0 | 2.3E-03 | 4.6E-03 |
| L*G | Rhizobium sp. | 617.5 | -3.3 | 1.0 | -3.2 | 1.2E-03 | 1.9E-03 |
| L | Saccharimonadota | 491.9 | 3.1 | 0.7 | 4.2 | 3.3E-05 | 4.4E-05 |
| B | Saccharimonadota | 491.9 | 2.4 | 0.7 | 3.3 | 1.0E-03 | 1.6E-03 |
| G | Saccharimonadota | 491.9 | 2.9 | 0.7 | 3.9 | 1.0E-04 | 2.0E-04 |

*Table S44:* Active microbial community differentially abundant taxa. First, twenty ASVs with the highest species scores on CAP 1 and CAP 2 from the CAP ordination of Bray-Curtis dissimilarities (distance matrix~ Treatment +block) were selected. These ASVs were aggregated to the lowest taxonomic level, typically the genus level. Then DESeq2 was run on these key taxa (rarefied reads ~ legume * grass * brassica). Model terms are the binary variables of the present/absence of a plant functional group in a plant mixture. The base mean (in rarefied reads) and log fold change are calculated from the plant-free soil pots. Only significant taxa are shown (p adj < 0.01).

| Model term | Lowest taxonomic ID | Base Mean | log2 Fold Change | lfcSE | Wald test stat | P value | P adj |
|---|---|---|---|---|---|---|---|
| L*G | Actinomycetes | 7018.4 | 3.6 | 1.2 | 3.0 | 2.3E-03 | 8.2E-03 |
| G | Actinomycetes | 7018.4 | -4.0 | 0.9 | -4.7 | 2.4E-06 | 8.5E-06 |
| B | Actinomycetes | 7018.4 | -3.7 | 0.9 | -4.2 | 2.9E-05 | 1.0E-04 |
| L | Actinomycetes | 7018.4 | -3.7 | 0.9 | -4.1 | 3.5E-05 | 2.5E-04 |
| L*G | Cyanobacterium | 318.6 | -7.6 | 1.9 | -4.0 | 7.4E-05 | 5.2E-04 |
| G | Cyanobacterium | 318.6 | 7.1 | 1.4 | 5.1 | 2.9E-07 | 2.0E-06 |
| B | Deinococcus sp. | 2393.7 | 9.0 | 1.8 | 5.1 | 4.1E-07 | 2.9E-06 |
| B | Escherichia sp. | 1165.0 | -5.4 | 1.5 | -3.6 | 3.6E-04 | 8.3E-04 |
